# The historical ecological background of West Nile virus in Portugal provides One Health opportunities into the future

**DOI:** 10.1101/2023.11.30.569416

**Authors:** Martim Afonso Geraldes, Mónica V. Cunha, Carlos Godinho, Ricardo Faustino de Lima, Marta Giovanetti, José Lourenço

**Affiliations:** Centre for Ecology, Evolution and Environmental Changes (cE3c), Faculdade de Ciências, Universidade de Lisboa, Lisboa, Portugal; Global Change and Sustainability Institute (CHANGE), Faculdade de Ciências, Universidade de Lisboa, Lisboa, Portugal; Biosystems & Integrative Sciences Institute (BioISI), Faculdade de Ciências, Universidade de Lisboa, Lisboa, Portugal; Universidade de Évora, Évora, Portugal; Faculdade de Ciências, Universidade de Lisboa, Lisboa, Portugal; Centro de Biodiversidade do Golfo da Guiné (CBGG), São Tomé, São Tomé and Príncipe; Laboratório de Flavivírus, Instituto Oswaldo Cruz, Fundação Oswaldo Cruz, Rio de Janeiro, Brazil; Instituto Rene Rachou, Fundação Oswaldo Cruz, Belo Horizonte, Minas Gerais, Brazil; Department of Science and Technology for Humans and the Environment, Università of Campus Bio-Medico di Roma, Italy; Climate amplified diseases and epidemics (CLIMADE) Americas, Brazil; Universidade Católica Portuguesa, Faculdade de Medicina, Biomedical research centre, Oeiras, Portugal; Climate amplified diseases and epidemics (CLIMADE) Europe, Portugal; Biosystems and Integrative Sciences Institute (BioISI), Faculdade de Ciências da Universidade de Lisboa Lisboa, Portugal

## Abstract

West Nile (WNV) is a zoonotic arbovirus with an expanding geographical range and epidemic activity in Europe. Not having yet experienced a human-associated epidemic, Portugal remains an outlier in the Mediterranean basin. In this study, we apply ecological niche modelling informed by WNV historical evidence (1969-2022) and a multitude of environmental variables from across Portugal. We identify that ecological backgrounds compatible with WNV historical circulation are mostly restricted to the south, characterized by a warmer and drier climate, high avian diversity, specific avian species and land types. We estimate WNV ecological suitability across the country, identifying overlaps with the distributions of the three relevant hosts (humans, birds, equines) for public and animal health. From this, we propose a category-based spatial framework providing first of a kind valuable insights for future WNV surveillance under the One Health nexus. We also forecast that climate trends alone will contribute to pushing adequate WNV ecological suitability northwards, toward regions with higher human density. This unique perspective on the past, present and future ecology of WNV addresses existing national knowledge gaps, enhances our understanding of the evolving emergence of WNV, and offers opportunities to prepare and respond to the first human-associated epidemic in Portugal.

## Introduction

West Nile virus (WNV) is a mosquito-borne virus of the *Flaviviridae* family, first identified in 1937 in the West Nile district of Uganda. The ecology of the virus is characterized by a zoonotic transmission cycle maintained between mosquitoes and avian species. Six mosquito genera have been implicated in transmission, but *Culex spp*. (e.g. *Culex pipiens*, *Culex univitattus / perexiguus, Culex modestus*) are commonly referred to as the main vectors in Europe and North America. WNV is genetically diverse and currently up to nine lineages have been proposed ^1^. Occasional zoonotic viral spillover can occur from the avian-mosquito cycle to humans, as well as to domesticated and wild mammals ^2–4^. Most human-associated outbreaks across the globe have been attributed to viral lineages 1, 2 and 5 ^5^.

Among incidental mammal hosts, humans and equines feature predominantly in epidemiological data, with equines often serving as a sentinel species in many countries. Contrary to most avian species, mammals are inefficient amplifier hosts, and hence cannot establish mosquito-mammal transmission cycles ^4,6,7^. Approximately 80% of human infections are asymptomatic, while the rest may develop mild or severe disease of neuroinvasive nature with an associated mortality risk ^8,9^. There are currently no licensed vaccines nor antiviral treatments for humans ^4,10^, but four equine licensed vaccines are on the market ^10^.

Introduction and increased epidemic activity of mosquito-borne viruses is being promoted by globalized trends related to climate and human activity ^11–15^. Historically, particularly during the 20th century, reported epidemic activity of WNV was mostly associated with African countries and Israel ^16^. At the turn of the century, awareness of WNV changed globally when it was introduced to New York city, from where it later became endemic across the USA and Canada ^17^. In the following decade, WNV also seemingly became more active in Europe, Middle East and Russia ^14,16^. Currently, WNV known epidemiology exhibits significant spatiotemporal heterogeneities both within and between countries, the drivers of which are not well characterized. The most well described driver is local climate variation, with well demonstrated effects on mosquito and avian life-cycles and population dynamics, consequently influencing local WNV epidemic activity, dispersal and persistence ^4,18,19^.

In Europe, recent increasing epidemic activity has been reported in France, Italy, Greece, Hungary, Romania, Bulgaria, Serbia and Ukraine ^20,21^. The largest epidemic ever was recorded across the continent in 2018, with the total of 2,083 cases exceeding cumulative infections since 2010 ^22^. For the first time in 2020, Spain witnessed a large epidemic involving human infections and deaths, including in regions not previously known to be affected by WNV ^23,24^. More recently in 2022, the second highest ever yearly number of locally acquired cases was reported in Europe at 1133, and simultaneously the number of cases in Italy was the highest ever on record ^25^.

These recent trends, particularly in countries around the Mediterranean basin with similar ecological conditions, highlight Portugal as an outlier. Despite evidence of WNV circulation in Portugal since 1969 ^26^, the country has not experienced a human-associated WNV epidemic. For reasons not well understood, only four serologically confirmed human infections have been reported in the country ^27,28^. It is possible that the rarity of human cases is partially explained by underreporting, driven by the combination of a low symptomatic rate ^8^, lack of medical and veterinary awareness, and current passive surveillance systems. To date, it remains unclear if the virus circulates endemically or not, with a total of only two genome sequences generated to date being unable to support either scenario ^28^. We have previously shown that accumulated evidence is largely concentrated in the south of Portugal where the climate is Warm Temperate Mediterranean, contrasting to that of the north which is Temperate Mediterranean (Köppen-Geiger classifications) ^27,29^. However, other factors that are universally understood to modulate WNV epidemic activity have previously not been explored within the Portuguese context (e.g. land types, host distributions).

In this study, we model and explore the ecological background and suitability for WNV transmission in Portugal, as informed by an historical dataset (1969-2022) of past evidence across the country and a large number of environmental variables. We identify that ecological backgrounds showing higher suitability for circulation (i.e compatible with evidence) are mostly restricted to the south of the country, generally defined by warmer temperatures, lower precipitation, high avian diversity, unique land types and prevalence of specific avian species. We leverage the ecological background and suitability outputs to propose a novel suitability and evidence based framework. We characterize regional potentials for future surveillance initiatives under a One Health perspective, including the three-host axes of WNV public and animal health importance (humans, equines and birds). Finally, given the identified, predominant role of local climate in our analyses, we perform a projection exercise accounting for observed, long-term climate change trends to estimate the changing landscape of ecological suitability across the country. We find that climate change alone will contribute to an expansion of regions suitable for WNV circulation. Together, our results reiterate that the south of Portugal is ecologically suitable for WNV transmission, and support a change towards active surveillance in the coming years.

## Results

We collected evidence for WNV in mainland Portugal (1969-2022). Given the existing passive surveillance framework and the fact that a large portion of the reported evidence did not include geographic coordinates, we summarized existing evidence by attributing present or pseudo-absent status per the smallest administrative geographical unit (*freguesia*, N=2882) (**Figure 1A**). Historical evidence included samples from birds, equines, humans, other mammals and vectors (**Figure 1B**). The predominant diagnostic methods were enzyme-linked immunosorbent assays (ELISA), viral neutralization testing (VNT), hemagglutination inhibition testing (HI), other serology-based methods, polymerase chain reaction (PCR), and viral isolation (**Figure 1C**).

**Figure 1.**
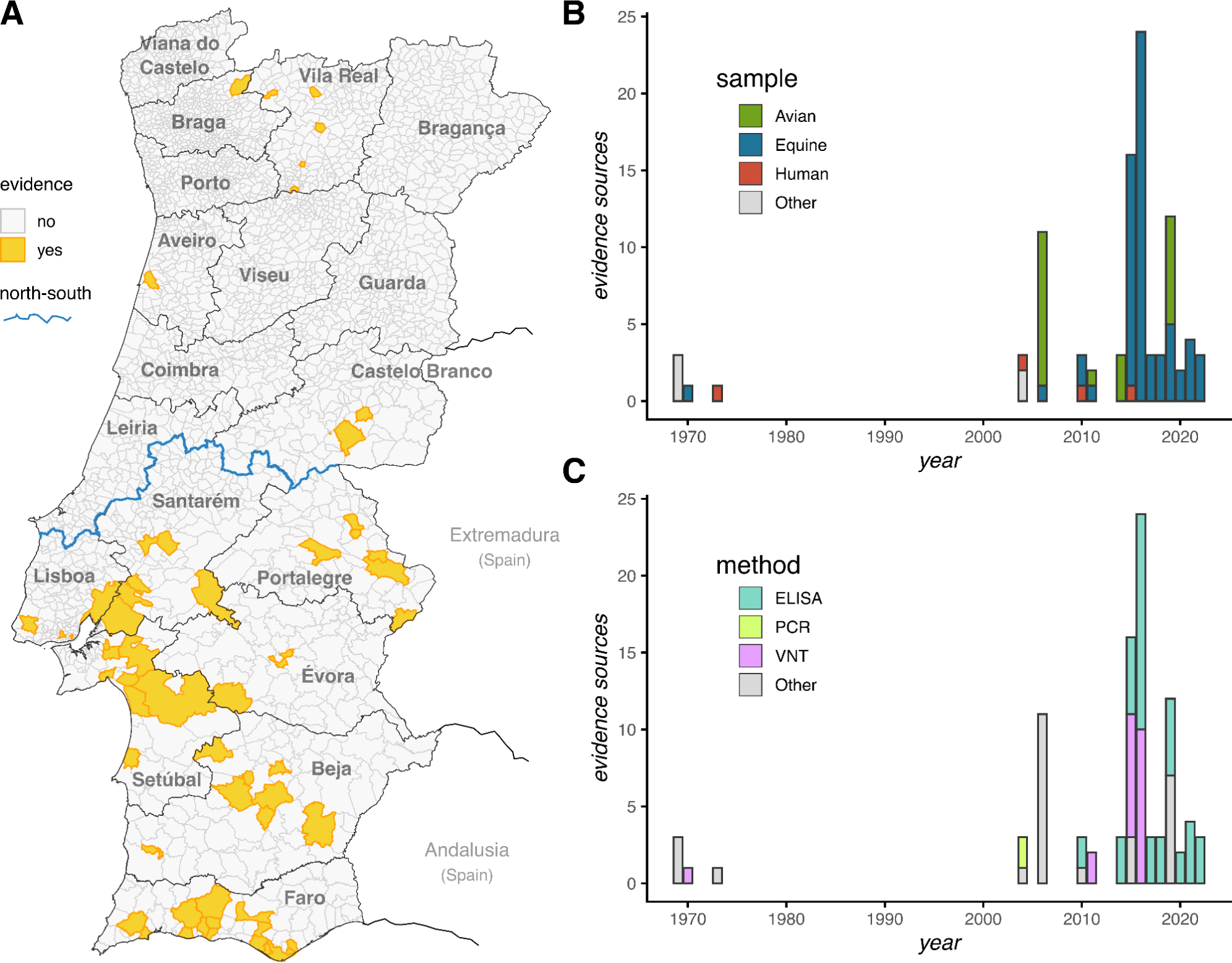
WNV historical evidence in Portugal (1969-2022). **(A)** Map of Portugal, presenting the geographical distribution of historical evidence for WNV presence (yellow). *Freguesias*, the smallest geographical units (N=2882) have grey borders, while districts (N=18) have black borders and are named. The blue line marks the north-south geographical division. The regions of Extremadura and Andalusia of Spain are also highlighted bordering to the south of Portugal. Total number of evidence sources by **(B)** sampled host and (**C)** testing method by year (of publication). Full dataset available in Supplementary Table S1.

Evidence of WNV circulation became more frequent post 2010, with a significant proportion (67%) of reports from equines. The spatial distribution of evidence seemingly did not follow any similar pattern to that of the main WNV vectors in Portugal (*Culex spp*) nor that of equines, both of which are essentially present across the entire latitudinal range of the country (**Supplementary Figure S1**). Interestingly, most evidence was from southern districts, mirroring evidence in the bordering Spanish regions of Extremadura and Andalusia (**Figure 1A**), where WNV has recently caused outbreaks within a range of host-species ^30,31^.

### Predictive variables

We collected 51 variables that potentially influence local WNV transmission in Portugal (**Supplementary Tables S2-3**). These predictive variables served as input for ecological niche modelling towards explaining the historical geographical distribution of WNV evidence. For this, we designed an approach based on an ensemble model supported by the output of two machine learning models: a boosted regression trees (BRT) and a random forest (RF) model. In essence, each machine learning model was informed by the predictive variables to classify the WNV evidence status of each of the 2882 *freguesias* (present, pseudo-absent) under a cross-validation subsampling strategy. The resulting classification from both models was then used as predictive variables to inform an ensemble model on a final classification exercise.

Firstly, Recursive Feature Elimination (RFE) was used to exclude the least informative variables while minimizing cost to model performance. In summary, we provided both BRT and RF models with a decreasing number of predictive variables, iteratively removing the variable with least measured importance. Performance of both models was stable while removing many variables, decreasing only after removing 34 of the 51 available variables (**Figure 2A**). The ROC from the BRT model was the most sensitive, decreasing early on, after removal of 21 variables. Based on this sensitivity, we applied a conservative, heuristic approach that would stop RFE once the BRT ROC would fall by more than 1% of its initial range (when using all 51 predictor variables). The BRT ROC fell below such initial performance range after removing the variable *Turdus merula* (Common Blackbird). The retained 18 variables after the RFE process are detailed in **Table 1**.

**Figure 2.**
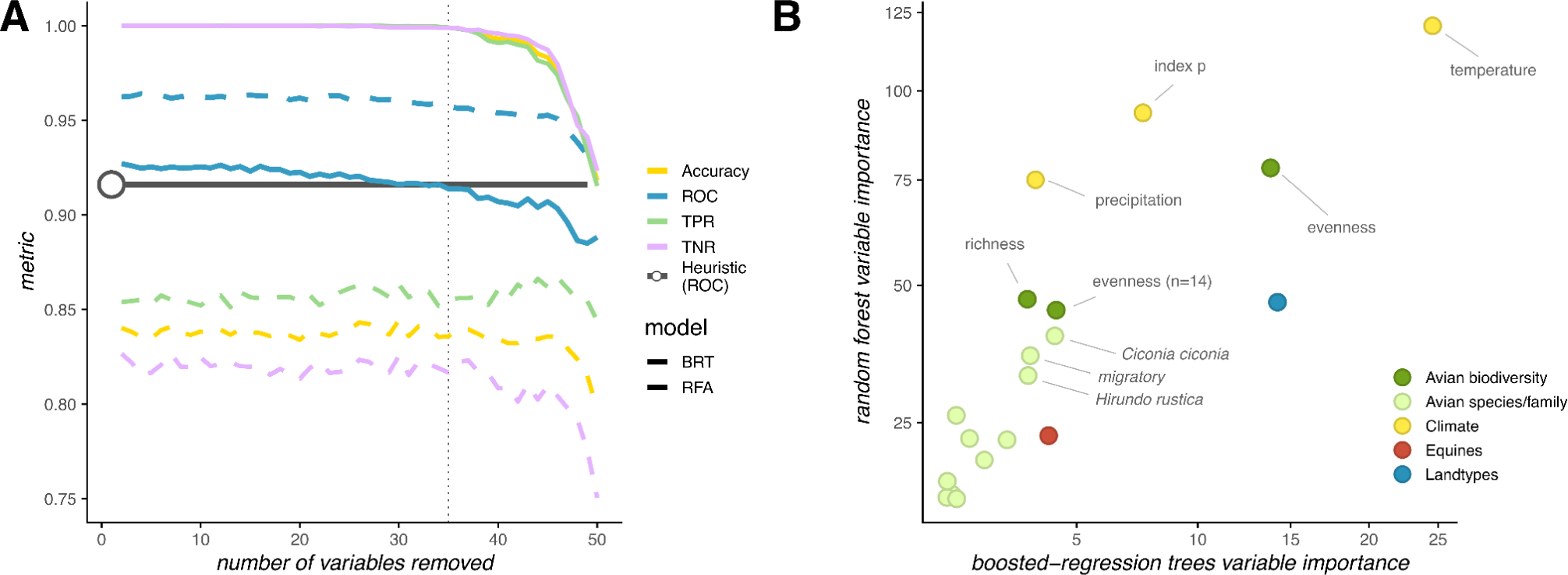
Metrics on recursive feature elimination and final variable importance. **(A)** Metrics of both the BRT (booster-regression trees, full lines) and RF (random forest, dashed lines) models when iteratively removing variables, one by one (ROC = Receiver Operating Characteristic; Accuracy = proportion of correct classifications; TPR = true positive rate or sensitivity defined as evidence status “present”; TNR = true negative rate or specificity defined as evidence status “pseudo-absent”). The dark grey line and point highlight the heuristic value used to stop the recursive feature elimination (1% decrease of the minimum of ROC BRT model, across the m-folds, when using all 51 variables). **(B)** Variable importance for both models using the final variables not excluded by the recursive feature elimination (N=18). Variables are aggregated into types (as in legend) for visualization and detailed in Table 1.

**Table 1.** Final set of 18 predictive variables from the Recursive Feature Elimination (RFE) approach. Sources, types and curation of variables detailed in Supplementary Table S2. * Relative to all other avian species.

| Variable | Description | Notes | Source |
| --- | --- | --- | --- |
| *Turdus merula* | Prevalence * | Common Blackbird | eBird <sup>47</sup> |
| *Garrulus glandarius* | Prevalence * | Eurasian Jay | eBird |
| *Hirundo rustica* | Prevalence * | Barn Swallow | eBird |
| *Erithacus rubecula* | Prevalence * | European Robin | eBird |
| *Anas platyrhynchos* | Prevalence * | Mallard | eBird |
| *Pica pica* | Prevalence * | Eurasian Magpie | eBird |
| *Columba palumbus* | Prevalence * | Wood Pigeon | eBird |
| *Ciconia ciconia* | Prevalence * | White Stork | eBird |
| *Cuculus canorus* | Prevalence * | Common Cuckoo | eBird |
| Corvids | Prevalence * | *Corvus corone*, *Corvus corax*, *Pica pica*, *Garrulus glandarius* | eBird |
| Migratory | Prevalence * | *Ciconia ciconia*, *Hirundo rustica* | eBird |
| Evenness (n=14) | Measure of prevalence balance | 14 species associated with WNV | eBird |
| Evenness | Measure of prevalence balance | All avian species | eBird |
| Equine | Prevalence | Absolute numbers | INE |
| Richness | Number of species | All avian species | eBird |
| Index P | Theoretical transmission potential | Climate-based | Literature <sup>27</sup> |
| Temperature | Absolute temperature | Mean Celsius | Copernicus.eu |
| Precipitation | Absolute precipitation in mm | Mean mm | Copernicus.eu |
| Land types | Most common land type | As classified in dataset | Copernicus.eu |

### Ecological niche modelling

Using the 18 selected variables by RFE (**Table 1**), we measured model performance in classifying WNV evidence status per administrative region (*freguesia*) (**Table 2**). The true positive rate (TPR) was higher than the true negative rate (TNR), and overall accuracy appeared capped by the performance of the TNR. On one hand this is an expected result from the highly diverse ecological background of *freguesias* with status pseudo-absent (covering the entire territory) compared to that of *freguesias* with status present (concentrated in the south). As such, it was more likely for the models to misclassify *freguesias* with pseudo-absent status given their diverse ecological backgrounds. On the other hand, it suggested that some *freguesias* with pseudo-absent status had similar ecological backgrounds to those with present status (i.e. predicting possible circulation in areas without existing evidence, explored in the results at a later stage).

**Table 2.** Final model performance using the selected 18 predictive variables. Results presented are for 200 m-folds. Sensitivity of each model’s performance for varying numbers of m-folds are presented in Supplementary Figure S2. ROC = Receiver Operating Characteristic; Accuracy = proportion of correct classifications; TPR = true positive rate (sensitivity) defined as evidence status “present”. TNR = true negative rate (specificity) defined as evidence status “pseudo-absent”.

| Model | ROC | Accuracy | TPR | TNR |
| --- | --- | --- | --- | --- |
| Random forest (RF) | 0.95 | 0.82 | 0.95 | 0.82 |
| Boosted-regression trees (BRT) | 0.94 | 0.81 | 0.93 | 0.81 |
| Ensemble (EN) | 0.95 | 0.82 | 0.94 | 0.82 |

#### Ecological background

The ecological background suitable for WNV circulation could be explored through the knowledge emerging from selecting predictive variables, their relative importance for the underlying models, and their relationships with the estimated likelihood of present status per *freguesia*.

The BRT and RF models generally agreed on the relative importance of the 18 predictive variables (**Supplementary Table S4**) with small variation, likely linked to their capacity to best capture linear (BRT) versus more complex (RF) variable relationships (**Figure 2B**). With highest importance were variables related to climate (temperature, index P, precipitation), landscape (land type) and avian biodiversity (evenness, calculated over all 288 avian species). Other avian biodiversity metrics were also selected but with less importance, such as biodiversity measured via richness (all species) and evenness over 14 hand-curated species of known interest for WNV transmission. The specific distributions of nine avian species were also deemed important although less than the climatic, landscape and most of the avian biodiversity measures. Finally, the distribution of equines was also selected but ranked low in importance compared to most other variables for both models.

Ecological backgrounds associated with WNV evidence status across the 2882 *freguesias* had properties in accordance with knowledge generated in previous studies. For example, ecological backgrounds with temperatures above 15 degree Celsius had a strong positive relationship with WNV presence (**Supplementary Text 1**) ^19,32^. In contrast, WNV presence was associated with backgrounds with lower precipitation, as also previously reported ^33,34^. The top land types associated with WNV presence included wetlands, shores (e.g. dunes), salt marshes, leisure facilities (e.g. golf courts), areas rich in agro-forest fields and estuaries ^35,36^. For measures related to the avian community, WNV presence was positively associated with higher biodiversity ^37^. For some specific avian species, WNV presence was simply dictated by the species presence or absence (e.g. *Ciconia ciconia, Erithacus rubecula, Passer domesticus*), while for others WNV presence had a monotonic relationship with prevalence ^37–39^. A full description on the relationships and contributions of all 18 variables to ecological suitability are included in **Supplementary Text 1**.

#### Ecological suitability

According to the ensemble model, different ecological backgrounds provided varying ecological suitability across the country (**Figure 3A**) (the independent output of the BRT and RF models is presented in **Supplementary Figure S3**, and all estimated suitability values per *freguesia* are made available in **Supplementary Table S5**). Highest suitability was estimated across the southern part of the country, namely across vast areas of the districts of Setúbal, Santarém, Portalegre, Évora, Beja and Faro, as well as more locally within the districts of Lisboa and Castelo Branco. The boundary of lowest to highest suitability between the north and the south colocalized with the separation of the two climate types present in the country (temperate mediterranean to the north, warm temperate mediterranean to the south, according to Köppen-Geiger classification ^29^). The identified southern region with high suitability was also in accordance with a recent study estimating risk of WNV transmission to humans and equines across the Iberian Peninsula using generalized linear modelling and a partially overlapping but different set of predictive variables (**Supplementary Figure S4**) ^30^. Notably, some smaller areas in the north were also estimated to harbour intermediate-to-high ecological suitability, namely around the urban centers of Aveiro, Vila Real and Covilhã, as well as the national park of Peneda-Gêres and the natural parks of Figueira de Castelo Rodrigo and Douro Internacional (**Figure 3A**).

**Figure 3.**
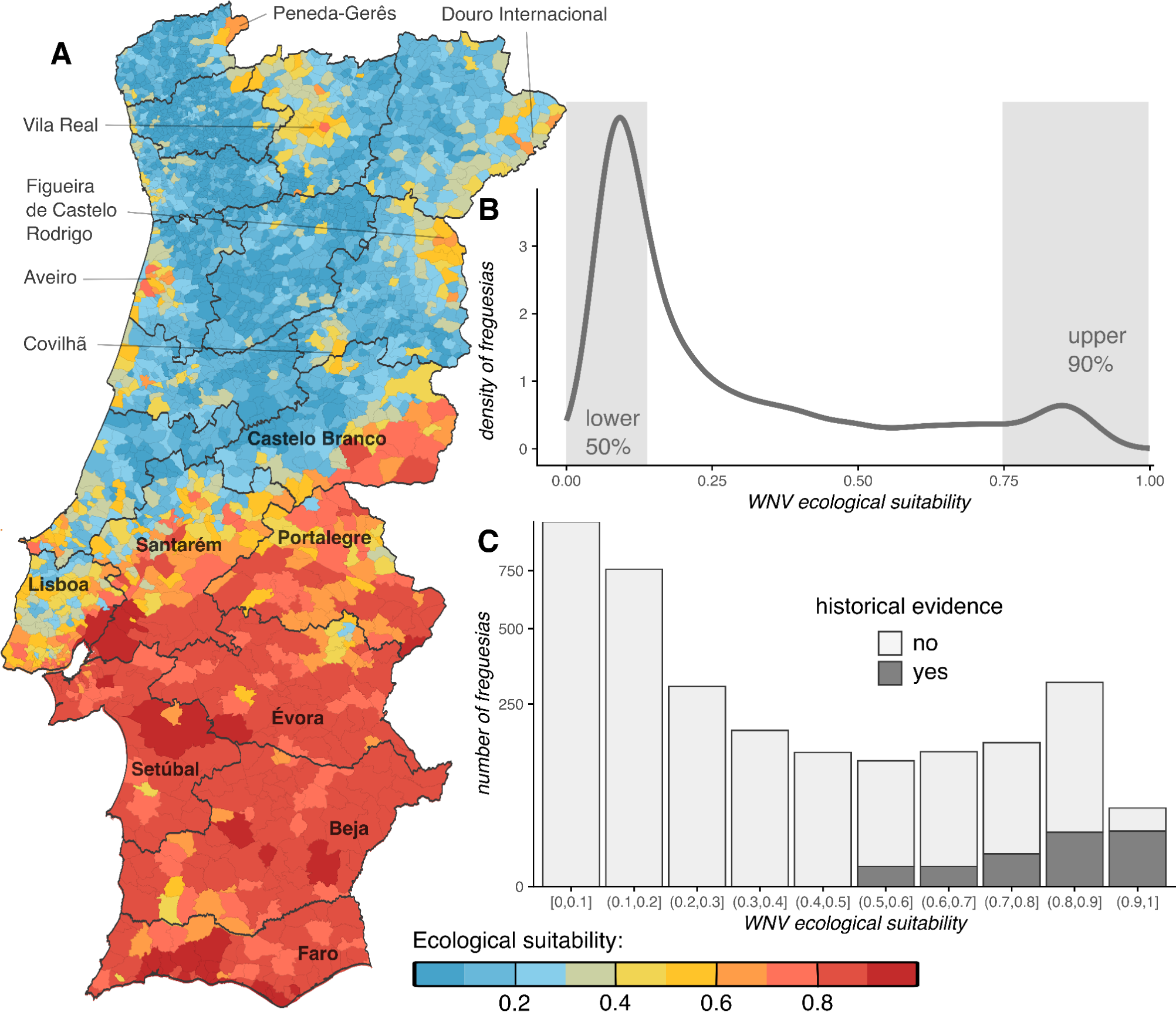
WNV estimated ecological suitability across Portugal. **(A)** Spatial distribution of estimated WNV ecological suitability per *freguesia* (color scale at the bottom). Dark grey spatial borders represent districts. **(B)** Density distribution of WNV ecological suitability across the country. The shaded area lower 50% represents the proportion of the distribution under the 50% percentile (lowest suitability), while upper 90% represents the proportion of the distribution above the 90% percentile (highest suitability). **(C)** Total number of *freguesias* with suitability within the bins of the x-axis. The number of *freguesias* with and without WNV historical evidence is presented by the different shades of grey according to the legend.

### Towards a One Health suitability and evidence based framework

The distribution of estimated ecological suitability across the country showed that *freguesias* with lowest suitability were dominant (**Figure 3B**) and at the same time lacked any historical evidence of WNV presence (**Figure 3C**). In contrast, *freguesias* with intermediate-to-highest suitability included freguesias with and without historical evidence (**Figure 3C**). The latter group highlighted the potential of the modelling approach to identify *freguesias* that, although lacking evidence, hold ecological backgrounds compatible with historical WNV circulation.

Proposing a novel suitability and evidence based framework, we defined three main categories (A to C) of epidemiological and public health importance based on different levels of estimated suitability. Categories related to intermediate and highest suitability were further divided into two sub-categories, depending on the existence of past evidence for WNV circulation (**Figure 4**). *Freguesias* with highest suitability distributed to the south (category A, **Figure 4A**) represented a broad spatial boundary in which future surveillance efforts, such as serological surveys of human and equine, molecular testing of mosquito and viral sequencing, are likely to provide valuable data. Within this boundary, those areas with past evidence of WNV circulation (sub-category A1, **Figure 4A**) represented subregions with greatest evidence-based support for future active surveillance. In contrast, areas with the main category C mainly distributed to the north (**Figure 4A**), had the lowest estimated suitability and lacked historical evidence of WNV. These areas highlighted the spatial boundary where active surveillance is least likely to provide new data, serving primarily to confirm the absence of WNV circulation. District-wise, Setúbal, Faro and Beja, all in the south, were identified as holding a dominance of *freguesias* classified with categories A and A1 within their territory (**Figure 4B**), thus presenting not just the highest estimated ecological suitability but also the greatest evidence for WNV circulation. The district of Évora, although presenting less evidence for circulation, was also highlighted as holding a dominance of *freguesias* with adequate ecological suitability across its territory. The main and sub-category of each of the 2882 *freguesias* are provided in **Supplementary Table S6**.

**Figure 4.**
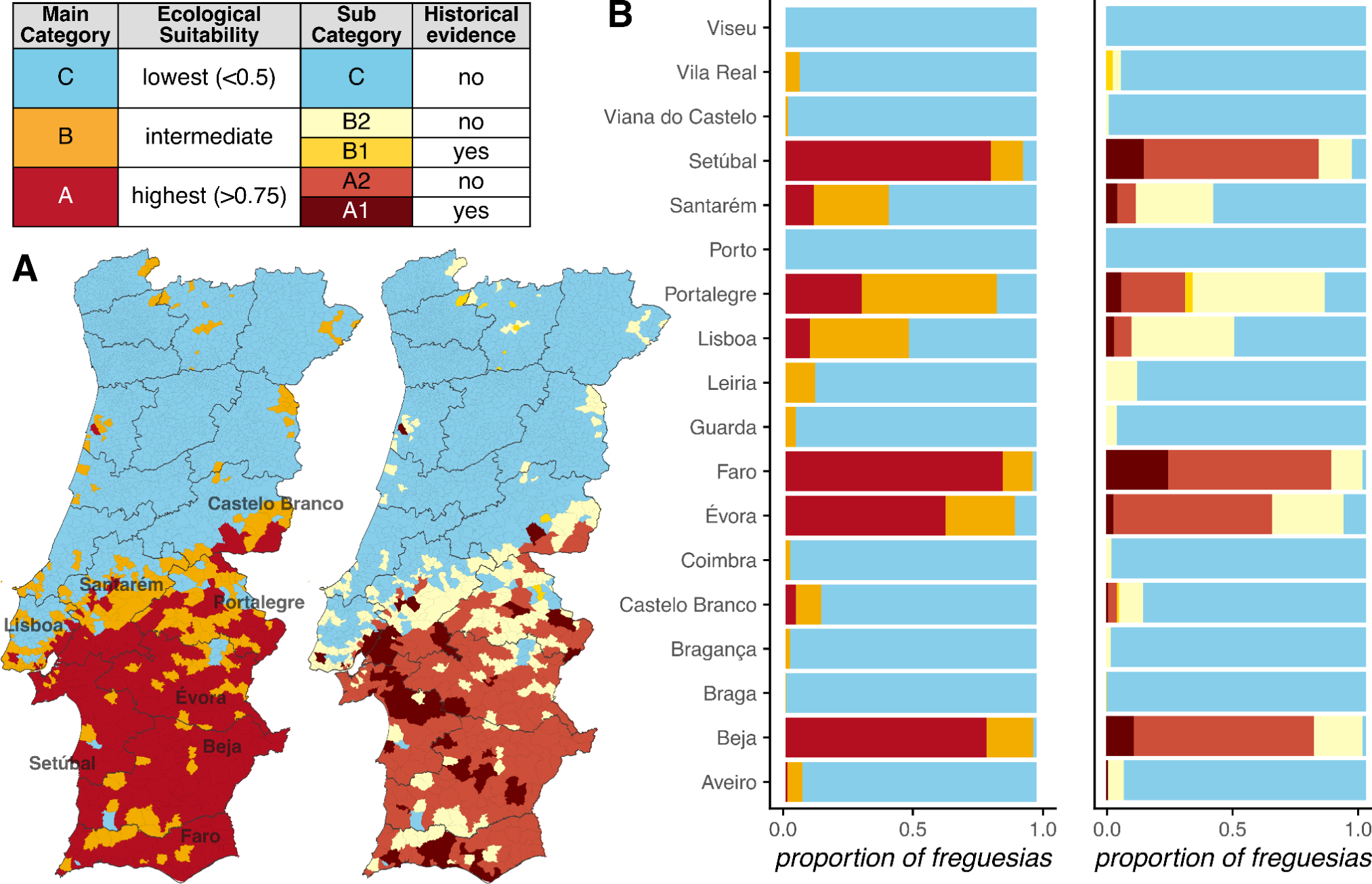
Proposed WNV ecological suitability and evidence based framework. **(A)** Spatial distribution of the three main proposed categories (left) and spatial distribution of the five proposed sub-categories (right). **(B)** Proportion of *freguesias* within each district that belong to the main (left) and sub-categories (right). **(A-B)** Both the maps and barplots are coloured according to the definitions of the categories as shown on the table (top left).

Towards a One Health perspective, we explored the spatial links between the suitability and evidence based main categories and the three-host axes of WNV epidemiology and public health – humans, equines and birds. In the main text we focused on identifying the *freguesias* that presented both the highest suitability (category A) and highest values of host distributions. A comprehensive bivariate visualization of all main categories and host ranges is presented in **Supplementary Figure 6**.

The Portuguese territory is highly heterogeneous in human population density, with the areas of highest density distributed along the western and southern coasts, with discrete hotspots in the interior **(Supplementary Figures S5-6**). Only *freguesias* in the south were identified as having both the highest suitability and human density (**Figure 5A**). The vast majority of identified *freguesias* belonged to the districts of Setúbal and Lisboa in the south-centre-west, and Faro in the south (**Figure 5B**). Critically, two of the three were districts in which all historical human WNV infections have been reported in the past (Setúbal, Faro).

**Figure 5.**
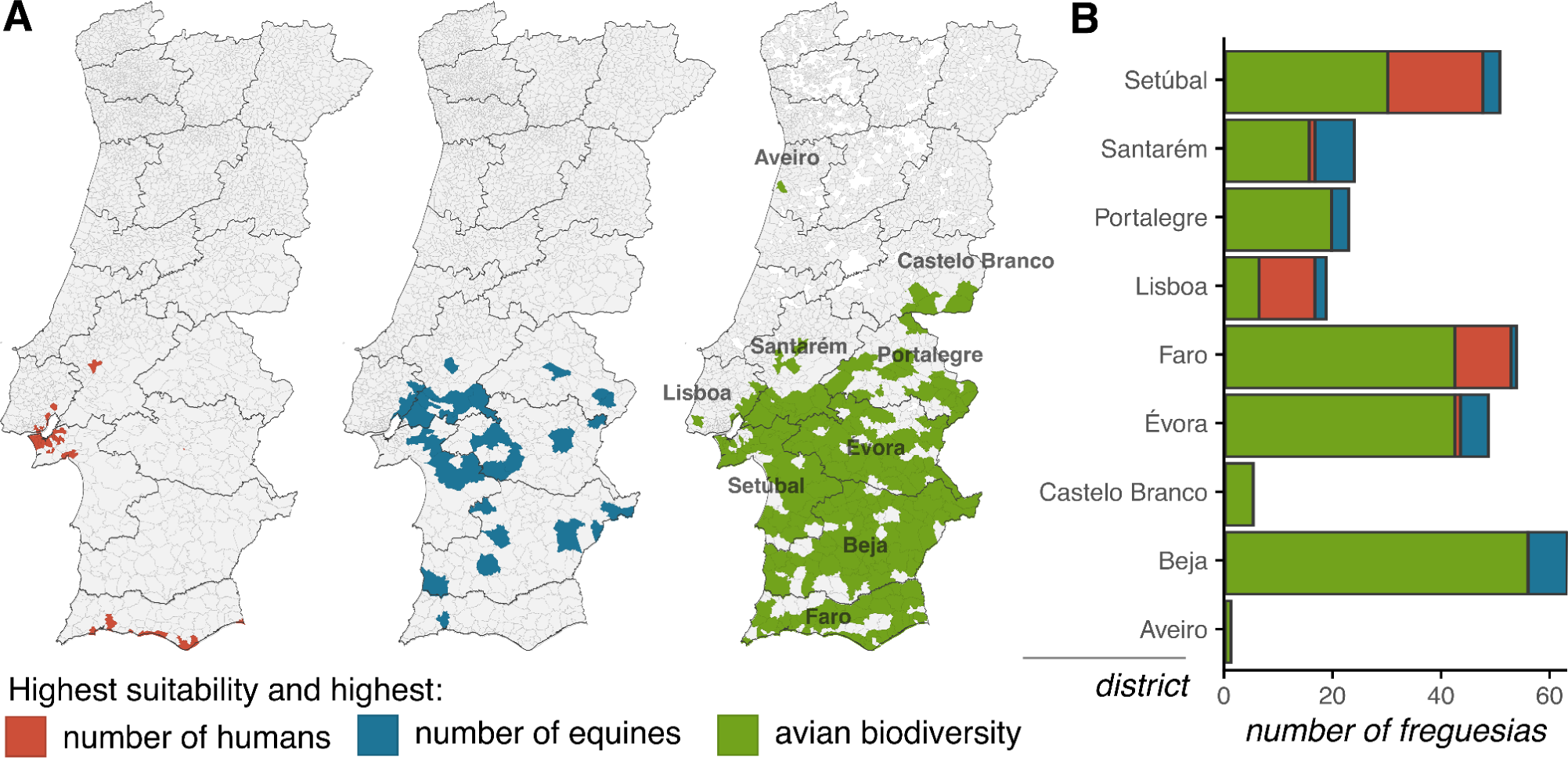
Overlap between WNV highest suitability and three-host axes. **(A)** Maps present the *freguesias* that hold both the highest ecological suitability (category A, Figure 4) and highest number of equines (left), number of humans (centre) and avian biodiversity (right, where white *freguesias* have missing data). **(B)** Summary of the total number of *freguesias*, per district, presented in panel A. Panels A and B use the color scale presented in the bottom.

In contrast to the distribution of humans, equines and avian biodiversity are less discrete across the country **(Supplementary Figures S5-6).** Overlap between category A and these non-human distributions revealed a number of relevant districts. The top 3 regarding the equine population were Beja, Évora and Santarém, located in the centre-south of the country. For avian biodiversity, the three most relevant also included Beja and Évora, as well as Faro, located in the centre-south and south, respectively. A full list of the identified districts and underlying *freguesias* in regards to their three-host background and the suitability and evidence based framework is included in **Supplementary Table S7.**

### Projecting the impact of climate change

Climate change has become increasingly evident in Portugal over the past decades. We have previously estimated that in the past ∼40 years (1981-2019) the country has become warmer and drier, slowly favouring the theoretical transmission potential of WNV ^27^. For example, by analysing satellite climate data for that period, we estimated a 0.04 degree Celsius yearly increase in mean temperature. Here, we applied the previously obtained estimates of climate change (Table 2 of ^27^) to the meteorological variables included in the ensemble model of this study (temperature, precipitation, index P), to project future changes to suitability up to the year 2050 (**Figure 6**).

**Figure 6.**
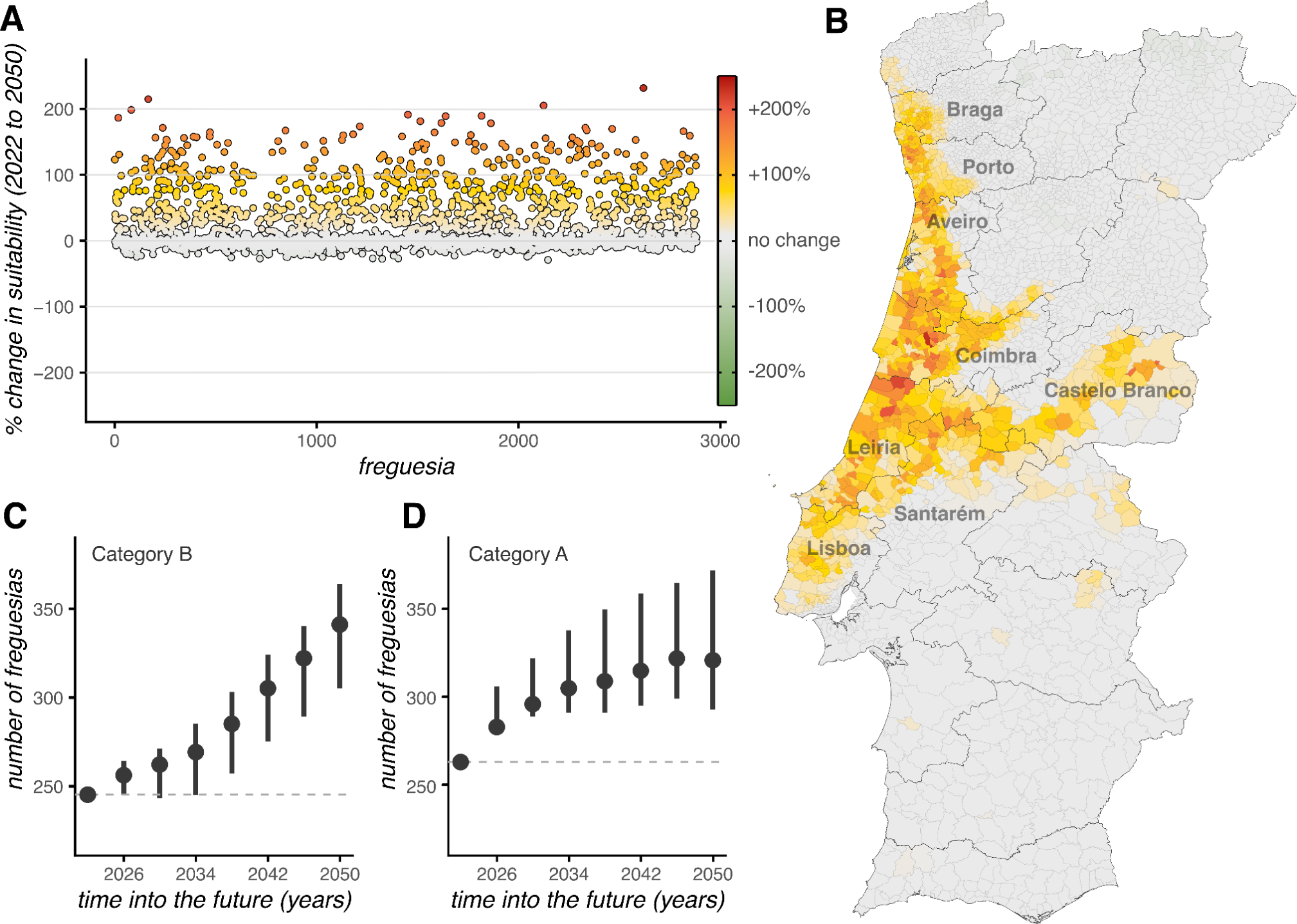
Contribution of climate change to suitability and evidence based categories. **(A)** Percent change in suitability per *freguesia* between the years 2022 and 2050. The x-axis presents all 2882 *freguesias* without any specific order. *Freguesias* (points) are colored according to whether suitability increased (warmer colors), remained mostly unchanged (grey), or decreased (colder colors). **(B)** Spatial distribution of *freguesias* presented in panel A. (A-B) Panels A and B share the color scale presented in panel A. **(C)** Change in time of the total number of *freguesias* holding category B as defined in Figure 4. **(D)** Change in time of the total number of *freguesias* holding category A as defined in Figure 4. In panels C-D, the whiskers represent the range of projections given the uncertainty in the rates of change of each climatic variable (see main text for source of rates).

We find that if climate trends remain similar to those of the past 40 years, WNV ecological suitability will remain stable across most of the territory, otherwise mostly increasing at specific locations (**Figure 6AB**). Overall, we estimate that approximately 62% of *freguesias* (N=1776) will experience almost no change (less than 10%) in the level of suitability they present in 2022 up to the year of 2050. Most change is expected in the north of the country in two fronts: one region across the west coast (Lisboa, Leiria, Coimbra, Aveiro, Porto and Braga), and another region across the north-south spatial boundary of the currently suitable south (Santarém, Castelo Branco) (**Figure 6B**). Approximately 6% of *freguesias* (N=179) will experience decreases in suitability, while 32% (N=927) will experience an increase. Decreases, however, are estimated to be only marginal in intensity when compared to increases (**Figure 6A**). Among *freguesias* with estimated increases by the year 2050, suitability will increase on average by 66% above the current values, but critically, 194 of these will experience more than a doubling (+100%) of their current suitability (**Figure 6AB**).

Following these trends, climate change will result in some *freguesias* changing from category C (lowest suitability) to category B (intermediate suitability), and others from category B to category A (highest suitability). Both the total number of *freguesias* with categories B and A will grow over the next decade (**Figure 6C-D**). Those with category B will grow in number from 245 in 2022 to 341 (range 305-364) in the year 2050 (**Figure 6C**), while those with category A will grow from 263 in 2022 to 321 (range 293-372) (**Figure 6D**). The vast majority of changes in main categories (C to B, and B to A) will take place in *freguesias* standing on the spatial boundaries of the current north-south divide in suitability (**Supplementary Figure S7**).

## Discussion

Portugal, standing on the far west of Europe, remains an outlier among its Mediterranean peers on not having yet experienced a WNV epidemic associated with human populations. To date, only four human cases have been reported, even with accumulating evidence that WNV circulates in the country and regularly crosses over its avian-centred transmission cycle into equine populations. Expanding previous attempts to decipher the uniqueness of the Portuguese context, we here apply ecological niche modelling based on machine learning techniques informed by an historical WNV evidence dataset (1969-2022) and a large number of environmental variables (N=51) from across the smallest administrative geographical units of the country (N=2882).

Leveraging on the modelling outputs, we show that ecological backgrounds compatible with past evidence for WNV circulation are mostly restricted to the south of the country, generally defined by a warmer and drier climate, the presence of higher avian diversity and a combination of unique avian species and land types. Overall, these results are in line with existing knowledge regarding the zoonotic cycle and spillover dynamics of WNV. For example, temperature is one of the most explored and validated factors driving the seasonality and spatial distribution of WNV incidence. In the context of reservoir hosts, we identify for the first time a group of specific species of national interest: Common Blackbird, Eurasian Jay, Barn Swallow, European Robin, Mallard, Eurasian Magpie, Common Wood Pigeon, White Stork, Common Cuckoo. Two groups of avian species, aggregated as migratory (White Stork, Barn Swallow) and belonging to the Corvid family (Carrion Crow, Northern Raven, Eurasian Magpie, Eurasian Jay) have also been identified as being of national interest. The fact that the distributions of specific species scored with lower importance than biodiversity measures also emphasized and reiterated that while individual species alone may contribute to local transmission, their contribution is limited in relation to the collective impact of the entire avian community. Similarly, while we noted the relevance of a few land types such as wetlands, the notable prominence of the generalized land type predictor, in contrast to any specific land type predictor, could be attributed to the known contribution of multiple land types to WNV transmission success.

Spatially, we also derived useful insights. The identified geographical, north-south boundary separating the lowest and highest suitability regions closely aligned with the demarcation of the existing temperate mediterranean (north) and warm mediterranean (south) climates. To the west, within the district of Lisboa where the capital city is, this boundary further coincided with the Tagus River, a national landmark for bird migration hotspots and avian biodiversity. Indeed, areas surrounding the capital were identified with high ecological suitability, going against the established assumption that, nationally, WNV favorable ecological conditions should be limited to the inner, rural and low human density areas of the country. We also identified regions with suitability that typically do not feature among those with past evidence for WNV circulation. For example, around the city of Aveiro, part of the Ria de Aveiro, an ecologically significant wetland, known for the high richness of nesting bird species. The localities of Figueira de Castelo Rodrigo, Peneda-Gerês and Douro Internacional, near protected natural parks, also showed up as suitable areas for WNV transmission. Finally, some of the identified localities with highest ecological suitability belonged to the districts of Setúbal (south-west) and Faro (furthest south). Among these were the localities where the only four human cases have been reported in Portugal. Ria Formosa in Faro, for example, the approximate location of two cases that occurred in birdwatchers, is an avian sanctuary that holds several resident and migratory species, making it a popular site for bird watching.

We considered 51 environmental variables, having selected 18 as relevant. While the latter allowed to define the historical ecological backgrounds of WNV, the exclusion of variables was informative on its own. For example, variables specific to human and other mammals (e.g. ovines, bovines) were excluded, validating the notion that transmission, wherever it has occurred, is not dependent on mammals, even if these animals can experience viral spillover. Of the climatic variables, only humidity was excluded, pointing to the fact temperature and precipitation hold sufficient information to capture the role of local climate. Variables specific to certain avian species were also excluded, including *Corvus corax* (common raven), *Turdus philomelos* (song thrush), *Parus major* (great tit), *Corvus corone* (carrion crow), *Passer domesticus* (house sparrow), suggesting low potential for future surveillance in these species in Portugal.

We estimated the direct effects of future climate trends to ecological suitability. In general, climate change alone will push the current north-south suitability geographical divide northwards, while keeping the south as the most suitable region. Climate change trends pushing the boundaries of mosquito-borne suitability northwards, specially in Europe and North America, is a topic of much ongoing research with a vast amount of supportive epidemiological and entomological evidence ^40^. Critically, the largest trends towards higher WNV suitability in Portugal up to the year 2050 will take place along the west coast where a large proportion of high density human populations. Climate change and demographics may thus coincidently raise the proportion of the human population at risk of WNV spillover into the near future.

An open question in Portugal pertains to whether the virus is endemic or epidemic (i.e. introduced annually or persisting over seasons). While we do not provide a definite answer, we generate sufficient output to propose that endemicity, if possible, would likely be favoured to the south. In fact, our dataset includes a noteworthy evidence point involving a symptomatic horse that tested positive for WNV antibodies off-season (on January 26 2022), in the *freguesia* of Comporta, Setúbal district. Our estimation of the ecological suitability for that *freguesia* is one of the highest in the country at 0.89. Together, this epidemiological event and our modelling output suggest potential, local overwintering of the virus. Support for endemicity could be tested in the future by the identification of unique, local viral lineages, but to date this has been not been feasible, with only two genome sequences available ^28^. From our proposed suitability and evidence based framework, we pinpoint districts (and their specific *freguesias*) where surveillance efforts would more likely result in positive molecular evidence and thus new isolates for sequencing. Sampling in animals could, for example, be performed within *freguesias* identified as sub-category A1 (high suitability with past evidence) and high equine density or high avian biodiversity, during typical months of WNV occurrence (May to October).

There are limitations to this work that should be revisited in future research once better and complementary data exists. Potentially, the largest of all is the predominance of equine-related evidence in the dataset. Although a consequence of the epidemiological history and passive surveillance in Portugal, it is possible that our outputs are biased to identifying ecological backgrounds that more specifically support spill-over events to equine populations. Nonetheless, it is reassuring that the equine variable was given far less modelling importance than climatic, land type and avian biodiversity variables, and that estimated ecological suitability was mostly restricted to the south in contrast to the equine distribution that included the entire north-south range. Other limitations include the use of variables measured from different time periods, a necessity given the scarcity of better sources for variables of interest. In terms of methods, we selected BRT and RF for being common in ecological studies and for having expertise with them. The RF showed a marginally better capacity in discriminating historical WNV presence-absence, which is consistent with other studies (e.g. ^41^). While the minor discrepancies in output between the models are expected, exploring alternative models, specially in light of new data, could be fruitful in the future. Regarding the projection of future climate effects, we did not include interactions with other factors. It is plausible that future climate trends will also affect other factors that were identified as drivers of suitability (e.g. distribution of avian biodiversity or the predominant local land type). As such, our results serve as a demonstration that climate change alone can alter and indeed expand the spatial boundaries suitable for WNV transmission, but the interaction with, and contribution of other factors should be addressed in future analyses.

In summary, our study contributes with a novel and unique perspective on the past, present and future ecology of WNV in Portugal, providing first of a kind valuable insights for decision-making into the future. From a One Health perspective, several districts are identified as holding adequate combinations of ecological suitability, past evidence and the presence of relevant hosts. Lisboa, in the west-centre region, for example, represents an almost virgin ground regarding WNV surveillance, but emerges as critical for future zoonotic potential into human populations. On the other hand, Faro in the far south, holds an optimal combination of factors that present opportunities to survey WNV current circulation, including in humans, equines and birds. Other districts such as Beja and Évora, instead, present opportunities for future studies focused mostly on non-human animals. The identified districts and underlying *freguesias* with varying relevance regarding the three-host axes form the initial basis of an information-driven framework under which future One Health surveillance should be based in Portugal.

Through efficient surveillance, geographical areas of risk can be identified and viral presence can be early detected, allowing for the implementation of timely interventions and mitigation measures both in humans and other animals. Active surveillance should include vector, human and other animal monitoring, as well as health education initiatives to raise veterinary, clinical and public awareness to WNV. Italy provides a good example of an European integrative One Health surveillance scheme with recent success ^42^. A shift towards a similar active surveillance with inclusion of genomic surveillance is recommended for Portugal in the near future. This shift can be guided by the suitability and evidence based framework proposed in this study. Only this way can we close existing gaps in knowledge, enhance our understanding of the evolving emergence of WNV, and be prepared to respond to the first and subsequent human-associated epidemics in the country.

## Methods

### WNV occurrence data

The evidence data for this study was collected from multiple sources to comprehensively assess the presence and distribution of WNV occurrences in Portugal. Initially, an occurrence list from our previous literature review was considered ^27^. To complement it, a partnership with Direção Geral de Alimentação e Veterinária (DGAV), the national veterinary and phytosanitary health authority was established, to include historical WNV outbreak information that was not publicly available. A second literature review was conducted to supplement the existing data up to 2022. Additionally, a recent master thesis by Costa et al.^43^ provided data on WNV serological testing of avian species. For analysis purposes, we summarized existing evidence into a binary classification of WNV “present” or “pseudo-absent” per the smallest geographical unit in Portugal, termed *freguesia*, of which there are 2882 in total. Regarding evidence based on avian samples, we considered only reports from known resident avian species, being thus conservative in considering, as much as possible, only evidence that pointed to avian infections within Portugal (rather than elsewhere and subsequently migrating to the country).

### Predictive variables data

A total of 51 biotic and abiotic environmental variables (here termed predictor variables) per *freguesia* were considered for analysis. These included factors related to climate, mammal and avian species, mosquitoes, land types, landscape, avian biodiversity and human population. Detailed information on variable selection, sources and curation is included in **Supplementary Text 1** and variable listing is included in **Supplementary Table S2**.

### Ecological niche modelling

We applied two machine learning (ML) approaches informed by the collected WNV occurrence data and predictive variables data: boosted regression trees (BRT) and random forests (RF). The output of these models was used to unravel and explore WNV ecological backgrounds (observed relationship of predictive variables with likelihood of WNV presence status) and quantifying WNV ecological suitability per *freguesia* (estimated likelihood of WNV presence status). The estimated WNV ecological suitability per *freguesia* of each ML model was combined to inform a metalearner model (Generalized Linear Model with negative weights) for a final estimation of local WNV ecological suitability per *freguesia*. For ML modelling classification of WNV present and pseudo-absent, we considered a conservative threshold of 0.5 (similar to previous studies ^44,45^). When performing machine learning with BRT and RF approaches, a balanced training dataset is recommended in terms of all possible classes for classification (in this case, present and pseudo-absent) ^46^. A total of 59 *freguesias* presented past evidence for WNV circulation. We thus devised a subsampling strategy, similar to k-folds in cross-validation, that involves multiple runs of the BRT and RF models (termed here m-folds), each including training subsets composed of 80 *freguesias* randomly sampled with replacement, with guaranteed equal proportions of *freguesias* with WNV present and pseudo-absent status. For modelling sensitivity and decision making, we considered up to 1000 m-folds. For some of the analyses (when explicitly mentioned), we determined an adequate, smaller than 1000 number of m-folds, by calculating and checking convergence of model performance (stopping at m-folds number above which more runs would not affect model performance). In effect, this approach guarantees that the smaller number of *freguesias* with WNV present status are compared and analyzed versus the larger number of *freguesias* with WNV pseudo-absent status, with modelling results presented as the mean output of the models across the m-folds runs, unless stated otherwise. A detailed description of the ML methods, parameterization and software packages used is included in **Supplementary Text 1**. All computational tasks were performed using R v4.2.1.

### Variable selection

We applied recursive feature selection by removing variables based on their measured importance in the BRT and RF models. After removal of the least important variable, the performance metrics of the models were recalculated, and the iterative removal of variables stopped once the most sensitive metric (identified in the results section as ROC from the BRT model) dropped by more than 1% of its minimum value across the m-folds when using all N=51 variables.

## Data Availability

All data is provided as **Supplementary Data File 1** (Excel), including WNV historical evidence (**Table S1**), details on the 51 predictive variables considered (**Table S2**), values of the 51 predictive variables considered per *freguesia* (**Table S3**), BRT and RF modelling variable importance (**Table S4**), estimated ecological suitability per *freguesia* (**Table S5**), categories per *freguesia* for the suitability and evidence based framework (**Table S6**), relevance status per *freguesia* related to the 3-host framework (**Table S7**).

## Supplementary Material

Supplementary material is included as **Supplementary Text 1** (Section 1 - Full description of data and methods; Section 2 - Details of the contribution of environmental variables to ecological suitability), **Supplementary Text 2** (supplementary Figures S1-6, and **Supplementary Data File 1** (Excel file, supplementary Tables S1-7).

## Acknowledgements

We thank Direção Geral de Alimentação e Veterinária (DGAV), the national veterinary and phytosanitary health authority, for engaging in this collaborative research project by providing valuable data on the historical occurrence of WNV in Portugal. We also thank José-María García-Carrasco and colleagues for providing the data holding modelling estimations of WNV risk in the Iberian Península ^30^, which was compared to our modelling outputs in Supplementary Material Text 1, working towards validation of both this and their study. We acknowledge the eBird platform, part of the European Breeding Bird Atlas 2 (EBBA2), for providing the distribution of avian species in mainland Portugal, as well as Colégio Tropical (CTROP) from the University of Lisbon for supporting a WNV research initiative in São Tomé linked to this research. Finally, we acknowledge strategic funding from Fundação para a Ciência e a Tecnologia (FCT, IP) to cE3c and BioISI Research Units (UIDB/00329/2020 and UIDB/04046/2020) and to the Associate Laboratory CHANGE (LA/P/0121/2020).

## Author Contributions

Conception and design: MAG, MVC and JL. Data collection: MAG, MVC, CG, RL and JL. Investigations: MAG, MVC and JL. Data Analysis: MAG, MVC and JL. Writing – Original: MAG, MVC and JL. Draft Preparation: MAG, MVC and JL. Revision: MAG, MVC, CG, RL, MG and JL. Resources: MVC, JL.

## Supplementary Material Text 1

### Section 1 Full description of data and methods

#### Data Sources and Curation

##### WNV occurrences data

In collaboration with a series of national partners we collected data from multiple sources to comprehensively assess the presence and distribution of WNV occurrences in Portugal. We started by considering our previous literature review on this topic, which missed key data collected by nacional entities ^1^. To complement it, we collaborated with *Direção Geral de Alimentação e Veterinária* (DGAV), the national veterinary and phytosanitary health authority, to obtain and include WNV outbreak information that wasn’t publicly available. Then, a second literature review was conducted to update on any new records available. We also obtained novel information on WNV serological testing of avian species from a recent master thesis conducted by Costa et al. ^2^.

Each evidence point did not necessarily correspond to a single evidence (e.g. single animal infection). In some instances, multiple animals tested positive at the same reported location, while in other cases, one point represented only a single evidence. To geographically represent the evidence points, latitude and longitude coordinates were utilized, with the freguesia assigned based on the location. In cases where the exact coordinates were not provided, the reported locality (name) was used to assign a freguesia. For mapping and further analysis purposes, we summarized existing evidence points into a binary classification system of WNV “present” or “pseudo-absent” per freguesia. Freguesias that reported sampling points were considered WNV “present”, and those with no past evidence were considered WNV “pseudo-absent”.

It should be noted that due to the passive nature of the existing surveillance system, in the vast majority of reported evidence, only positive events are reported. As such, the lack of reports in a freguesia does not necessarily imply true absence, but rather pseudo-absence due to potential underreporting. This transformation resulted in 59 freguesias demonstrating past evidence for WNV circulation. Such freguesias were mainly restricted to the districts of Faro, Beja, Setúbal, Évora, Lisboa, Portalegre, and Santarém. In contrast, 2823 freguesias lacked any documented record of WNV presence. Furthermore, in the context of avian evidence, we considered only occurrences from known resident species, being thus conservative in considering only evidence that pointed to infections within Portugal (rather than elsewhere and subsequently migrating to Portugal).

##### Predictive variables data

An extensive list of biotic and abiotic environmental variables (termed predictor variables) from Portugal, including climate, distribution of mammal, mosquito, and avian species, as well as landscape, biodiversity metrics and human-associated variables were used to inform the ensemble model. All variables are listed in detail in **Supplementary Table S2**.

Temperature, rainfall and humidity were considered. For each one, the average value for each freguesia from 1980-2021 was calculated using the Copernicus dataset “Essential climate variables for the assessment of climate variability” ^3^, which includes climatic variables at a time resolution of 1 month (1980–2021) and spatial resolution of 0.25°x 0.25° (25 km x 25km).

Topography-wise, we considered altitude and land types. Elevation is known to negatively affect *C. pipiens* abundance when higher than 1700m. Land type (e.g. rural, agricultural, forestry, wetlands and periurban) has previously been associated with mosquito abundance and WNV transmission ^4^. Elevation data was downloaded from the EU-DEM v1.1 dataset, which has a resolution of 25m x 25m ^5^. The average elevation per freguesias was subsequently used. The land type dataset used was “Land cover classification gridded maps from 1992 to present derived from satellite observations” which contains 44 different land classification classes for 1990, 2000, 2006, 2012 and 2018 with a resolution of 100m x 100m ^6^. These classes were aggregated into six categories of interest in the context of WNV: water bodies and courses (classes 511 and 512), forests (classes 311, 312, 313), rice fields (class 213), pastures (class 231), road and rail networks and similar land types (class 112). The proportion of land covered by each land type category per freguesia was then calculated. Finally, a freguesia-level variable was created containing the most common land type (mode), between the 44 classes, for each freguesia. Finally, we also aggregated three land type classes: natural (classes 311, 312, 313, 321, 322, 323, 324, 331, 332, 333, 334, 335, 411, 412, 421, 422, 423, 511, 512, 521, 522, 523), agroforest (classes 211, 212, 213, 221, 222, 223, 231, 241, 242, 243, 244) and urban (classes 111, 112, 121, 122, 123, 124, 131, 132, 133, 141, 142). This aggregation was used to create a categorical predictor called “land3classes” that identifies the predominant land type of each freguesia (between rural, agroforest and urban).

We also considered a climate-based suitability index, termed Index P, which measures the theoretical transmission potential of WNV as dictated by temperature and humidity. The index has been demonstrated to be correlated with spatiotemporal WNV incidence data of humans and equines, e.g. in Israel REF. We used the spatial dataset made available in the 2022 Portuguese study to calculate a mean value of index P per freguesia ^1^.

To represent the WNV animal reservoir, we used the distribution of several avian species made available on the eBird platform ^7^. This is the most common based online database for bird records in Portugal, aggregating, in addition to voluntary records, various monitoring schemes on a national scale. Previous to data analyses, we removed eBird records not suitable for this research (e.g. sea counts), records impossible to associate with a freguesia, and duplicated records (eBird allows for multiple observers to share the same record list). We used data from 2015-2021 which includes records between March and May. The relative frequency of 288 species as observed per freguesia was used, and we further estimated biodiversity metrics such as species richness, evenness, Shannon Index and Simpson Index, using the R package vegan v2.6 ^8^. To avoid modelling the 288 available species, we defined high-priority species under three categories as in Vaughan et al ^9^: (1) birds known to produce high WNV viremia; (2) birds that are typically numerous and associated with humans; (3) and birds that are preferentially fed on by mosquitoes of importance for WNV transmission. For (1) we selected *Corvus corax, Corvus corone* and *Garrulus glandarius*. In category (2) *Erithacus rubecula, Turdus merula, Turdus philomelos* and *Pica pica* were selected. *Columba palumbus, Passer domesticus* and *Anas platyrhynchos* were taken to represent category (3). Portugal is in a migratory bird flyway, so we further selected two migratory species common in Portugal during migration season that have both tested positive for WNV: White Stork (*Ciconia ciconia*) and the Barn swallow (*Hirundo rustica*). The common Cuckoo (*Cuculus canorus*) and Great Tit (*Parus major*) are both abundant in Portugal and have been studied for WNV seroprevalence in Spain ^10^. Finally, we further aggregated *Corvus corone, Corvus corax, Pica pica* and *Garrulus glandarius* in a category named *corvids*, and *Ciconia Ciconia* and *Hirundo rustica* in a separate category called *migratory*. In the main text we also refer to avian species by their English common name, defined according IOC V13.1 (https://www.worldbirdnames.org/new/).

Several dead-end hosts (mainly mammals) have been demonstrated to test positive for WNV infection. In addition, the presence of ovine, bovine and caprine farms within a 500 m radius from a mosquito trapping point has been considered a positive predictor for *Culex pipiens* abundance ^4^. We thus considered the spatial distribution of several dead-end mammal hosts such as equine, caprine, bovine, swine, ovine, poultry and rabbits. Data was sourced from the National Institute of Statistics (INE), officially called “Efectivo animal (N.o) das explorações agrícolas por Localização geográfica (NUTS - 2013) e Espécie animal (Modo de produção biológico); Decenal” ^11^. The data consists of two datasets containing the number of each mentioned group per freguesia for 2009 and 2019. The average between those years was calculated per freguesia.

The Human footprint index (HFI) – an index of human pressure, incorporating built environments, energy and transportation infrastructure, agricultural lands, and human population density – was previously found to be an important predictor of vector-borne disease occurrence ^12^. We downloaded HFI from 2000 to 2013 and averaged it by freguesia.

Human population density was calculated per freguesia using the INE nationwide census of 2021 ^13^.

#### Ecological niche modelling

All computational tasks were performed using R V4.2.1 utilizing packages gbm V2.8.2.1 ^14^, caret V6.0-93^15^, pROC V1.18.0 ^15,16^, h2o V3.42 (for BRT and RF models) ^17^.

##### Modelling framework

When performing machine learning with BRT and RF approaches, a balanced training dataset is recommended in terms of data entries representing all possible classes for classification ^18^. To accomplish this, we devised a subsampling strategy, similar to k-folds in cross-validation, that involves iterative runs (m-folds) of the models with training subsets composed of 80 freguesias randomly sampled with replacement from the total 2,882 freguesias, with equal proportions of freguesias with WNV presence and pseudo-absence status. We considered a maximum of 1,000 m-folds, ensuring coverage of all freguesias across model runs and thus all possible ecological backgrounds (due to the vast number of possible sampling combinations, the likelihood of encountering the same training dataset in distinct runs was negligible). For some of the analyses (when mentioned in the maintext) we determined an adequate minimal number of m-folds, by calculating and checking convergence of model performance (stopping at m-folds number above which more runs would not affect model performance).

To explore ecological backgrounds (relationship of predictive variables with likelihood of WNV presence status) and quantifying the ecological suitability (likelihood of WNV presence status) of WNV, the m-fold strategy was employed with 200 m-folds (sensitivity presented in **Figure ST1**). The BRT model was configured with ntree equal to 50, lr of 0.1, and a tc of 0.01. The RF model was configured with ntree value of 50, with the maximum number of variables considered at each split (mtry) set to the square root of the total number of predictors in the dataset. Both models underwent resampling using a 5-fold cross-validation approach.

**Figure ST1.**
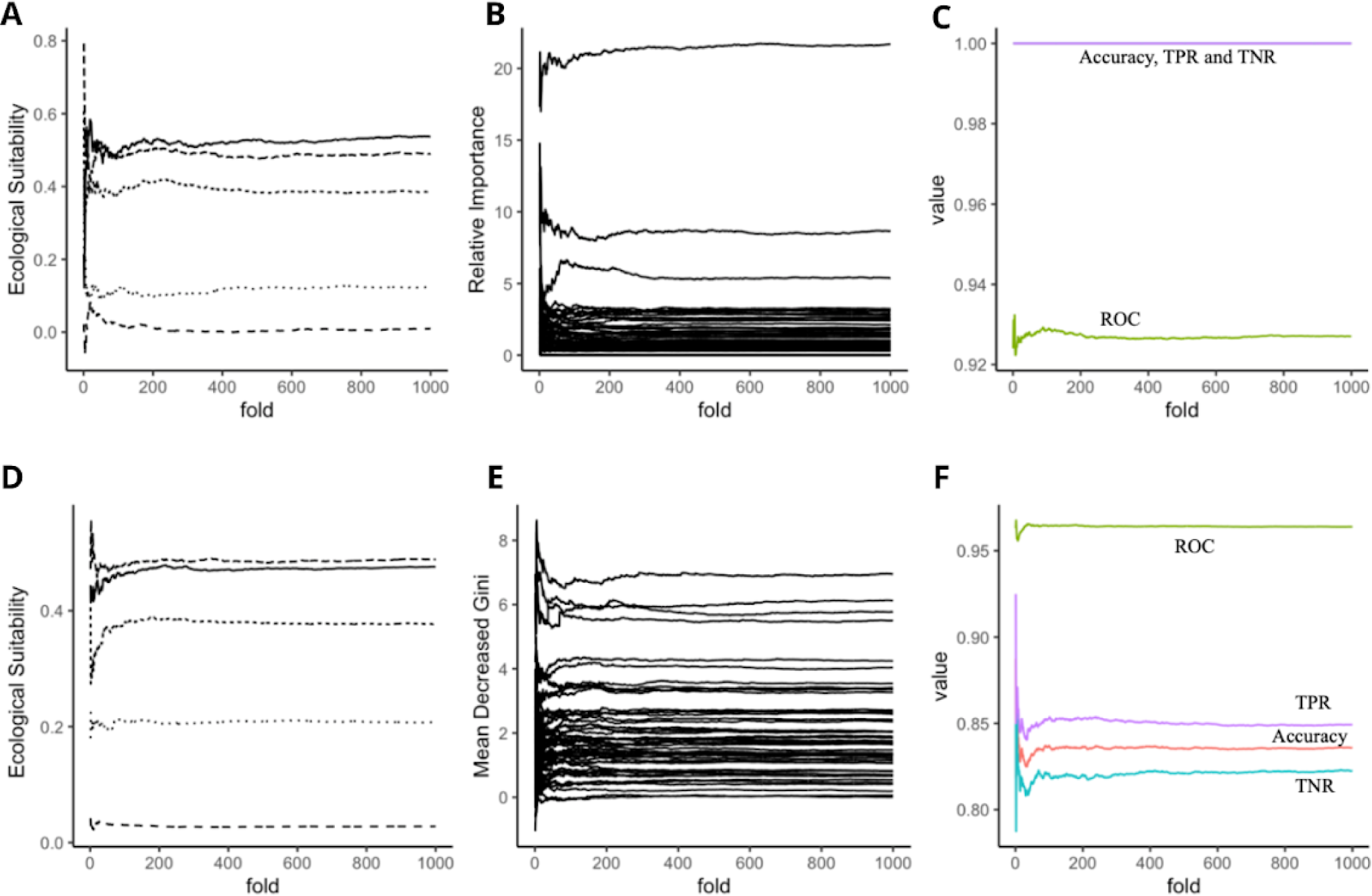
Sensitivity of model output to the variation considering different numbers of m-folds from 0 to 1000. WNV ecological suitability for five random freguesias (namely: União das freguesias de Cedofeita, Santo Ildefonso, Sé, Miragaia, São Nicolau e Vitória; Laundos; Penamacor; União das freguesias de Sameice e Santa Eulália and Tamanhos), for (A) BRT (boosted-regression trees) and (D) RF (random forest) models. Predictor variables’ importance for (B) BRT and (E) RF. Quality metrics: in green is represented ROC, in purple TPR (true positive rate), in red Aaccuracy, and in blue TNR (true negative rate) for (C) BRT and (F) RF models.

Finally, the predictions of both models were combined to inform a Super Learner known as Stacked Ensemble. We selected the metalearner model to be a Generalized Linear Model (GLM) with negative weights, a strategy that is becoming more and more frequent and has already been used in this field, for example, for modelling WNV probability in Florida ^19^.

We considered a conservative threshold of 0.5 (like in previous similar studies ^20,21^) for classification (i.e. pseudo-absent classification was considered below 0.5 and present classification above 0.5). We further conducted 10 repetitions of the ecological niche modelling to assess the degree of variability across different runs, and present the mean and variation of the ensemble model response in **Figure ST2.**

**Figure ST2.**
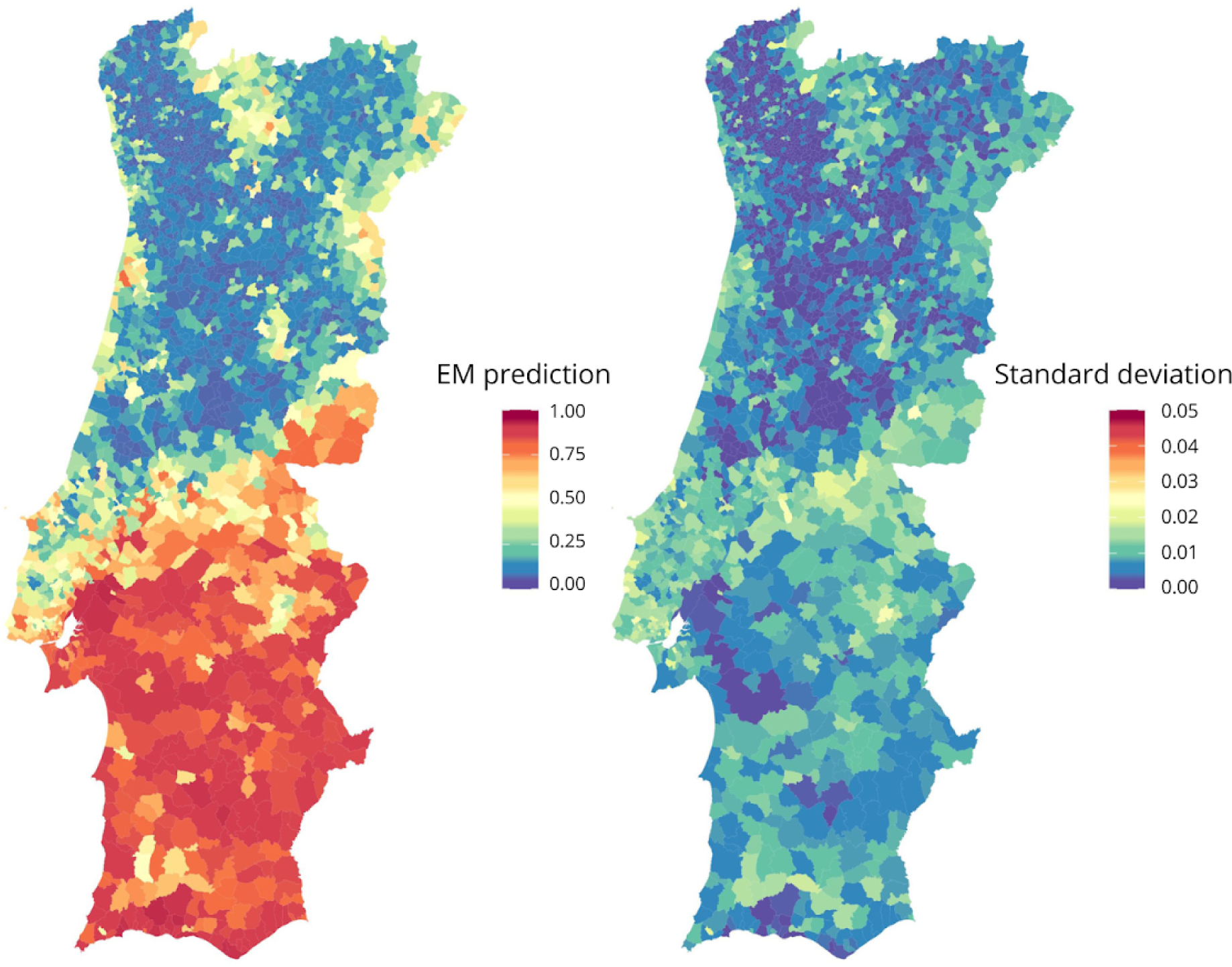
Final output of ecological modelling in terms of the spatial distribution of ecological suitability (ensemble model). (left) Mean ecological suitability and (right) Standard deviation of ecological suitability from model output. Maps show the N=2883 freguesias of Portugal.

##### Variable selection

Recursive feature elimination (RFE) is a prominent technique for discarding redundant (predictive) variables, retaining only a reduced set of variables that offer reproducibility and local scalability, as collecting a reduced set of variables for multiple contexts is more feasible ^22^. Typically, variables are iteratively removed based on their measured importance for the classification exercise. After removal of the least important variable, model performance metrics are recalculated, and the iterative removal of variables stops once a specific performance metric drops below a selected threshold. We measured variable importance for BRT and RF using relative importance (RI) and mean decreased Gini (MDG), respectively. When two different variables were identified as the least important in the BRT and RF models, both were removed. We used the performance metric ROC of the BRT model to decide when to stop removing variables (see details in main text). The BRT model employed a Gaussian loss function, with a total of 500 trees, a lr of 0.1, a tc of 1, and resampling performed using a 5-fold cross-validation approach. Conversely, the RF model employed the value of 500 trees and utilised a 10-fold cross-validation resampling method. The optimal value for mtry, which maximized accuracy, was selected based on the highest value achieved.

### Section 2 Details of the contribution of environmental variables to ecological suitability

The contribution of environmental variables to model predictions (ecological suitability) is here presented using partial dependence plots, to gain insights and even define quantitatively the ecological background of WNV presence-absence. Partial dependence plots were generated by contrasting the estimated ecological suitability (y-axis) versus the values of each variable per freguesia (x-axis); the latter discretized in several ranges for better visualization. The following text related to **Figures ST3** to **ST6** are based on the 18 predictor variables selected (out of the total 53) by the RFE approach described in the **main text**.

#### Estimated ecological suitability and climate-related factors

We observed a positive correlation between ecological suitability and temperature (**Figure ST3**). Mean, minimum and maximum daily, weekly, monthly and annual temperatures have been demonstrated to be positively related to the abundance of mosquitoes, although not linearly, since abundance may be reduced at too high temperatures ^4^ (not observed on the temperature range of Portugal).

**Figure ST3.**
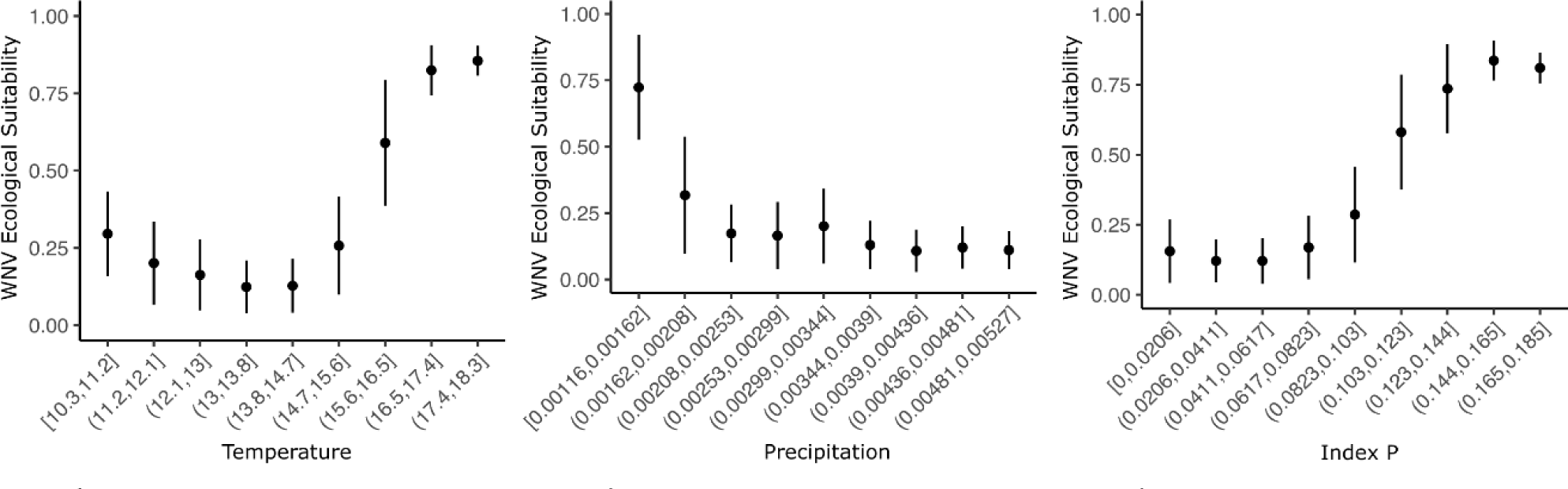
Partial plots of WNV ecological suitability in relation to climate. From left to right: temperature, precipitation and Index P. Temperature and Index P, as opposed to precipitation, reveal a positive relation with WNV ecological suitability. Variables are discretized into bins for a more intuitive visualization.

Similarly, Index P exhibited a positive relationship with suitability, aligning with its strong sensitivity to temperature fluctuations (**Figure ST3**) ^23,24^.

Conversely, precipitation displayed a negative association with suitability (**Figure ST3**), a trend commonly identified in existing WNV modelling studies ^25,26^. Explaining the nuanced role of precipitation as a driver for WNV circulation is challenging. On one hand, mosquitoes rely on water availability to lay eggs and complete their reproduction cycle. Therefore, by creating potential breeding sites, precipitation is believed to elevate vector abundance. However, in non-tropical countries like Portugal, precipitation exhibits a negative correlation with temperature. In such regions, precipitation typically coincides with low monthly temperatures, which may limit mosquito activity and subsequently impact WNV transmission dynamics. Positive relationships of WNV activity with precipitation are typically more easily found when considering lag periods between the two variables (e.g. ^27^).

#### Estimated ecological suitability and land type related variables

Land use and land cover are known to influence the distribution of WNV transmission cycle, vectors and reservoirs ^4^. *Culex pipiens’* abundance, for example, is commonly associated with rural, agricultural, forestal and peri-urban habitats. By analyzing each of the 44 land type classes available in our dataset, we could further explore the extent to which each one of them influenced the estimation of WNV ecological suitability in Portugal (**Figure ST4**). Leisure facilities appeared as the land type with the highest ecological suitability. This category includes, for example, football fields and golf courses whose irrigation creates good breeding sites for mosquitoes. The land types that followed in importance were tidal flats (wetlands), dunes/sands, salt marshes, agroforestry areas, green urban areas and estuaries, thus confirming the idea that areas that relate to shores, or somehow combine water sources with green areas are compatible with evidence of past circulation of WNV in Portugal. In its neighbor country Spain, the presence of wetlands in a 500 meter radius was identified recently as a positive predictor for WNV circulation ^4^. This happens because vectors can find sources for laying eggs, as well as birds that in their migrations rest in those areas, thus mixing vector and reservoir ^28^. In contrast, the land types least associated with adequate ecological suitability included permanent crops, airports and pasture fields.

**Figure ST4.**
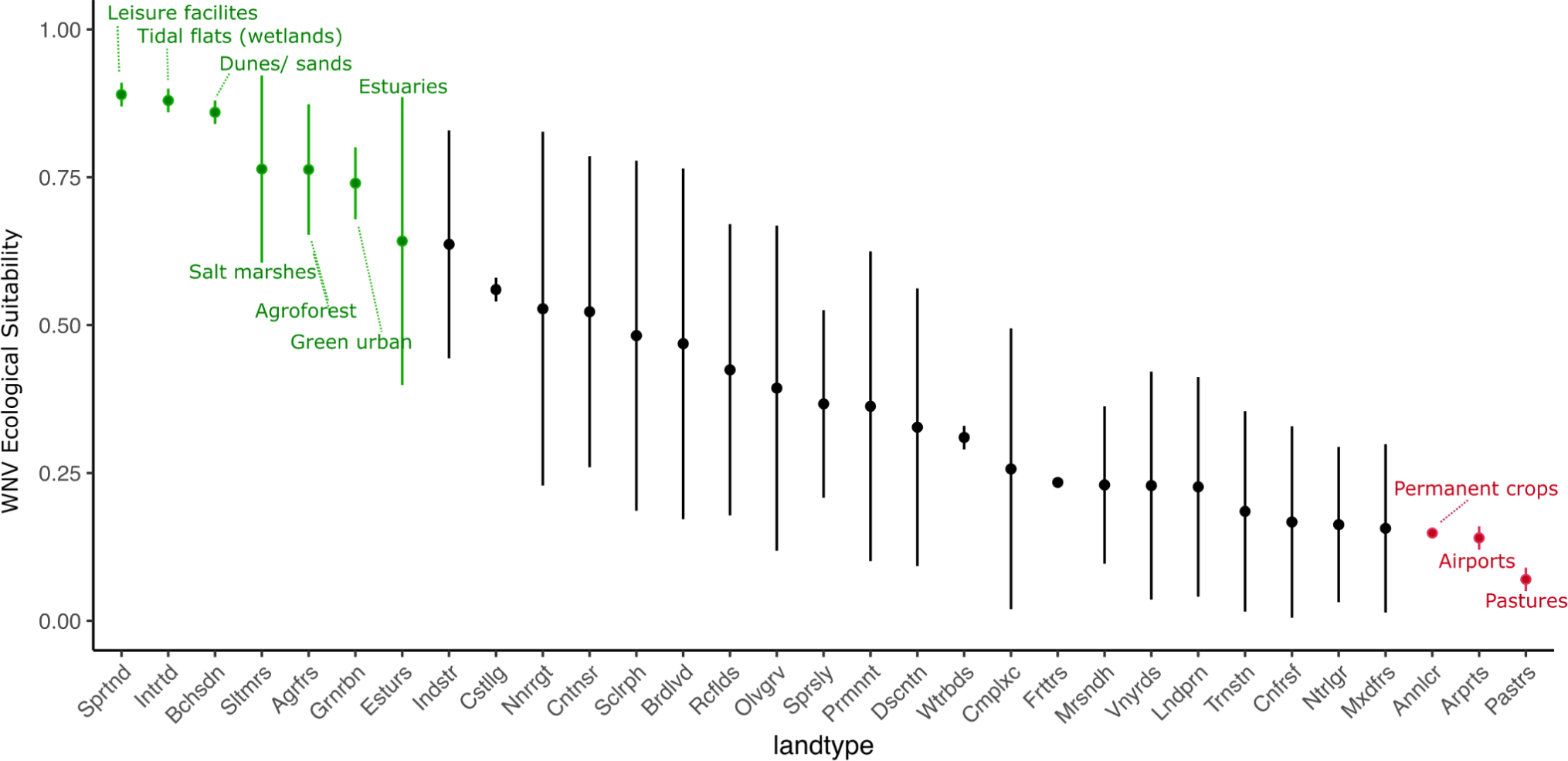
Partial plots of WNV ecological suitability in relation to land types. Partial plot demonstrating the average WNV ecological suitability from freguesias that have the given land type as the most common one. From left to right (highest to lowest ecological suitability), the land types are: Sports and leisure facilities; Intertidal flats; Beaches, dunes, sands; Salt marshes; Agroforestry areas; Green Urban Areas; Estuaries; Industrial or commercial units; Coastal lagoons; Continuous urban fabric; Non irrigated arable land; Sclerophyllous vegetation; Broad-leaved forest; Rice fields; Olive groves; Permanently irrigated land; Sparsely vegetated areas; Discontinuous urban fabric; Water bodies; Complex cultivation patterns; Vineyards; Fruit Trees and berry plantations; Moors and heathland; Land principally occupied by agriculture and significant areas of natural vegetation; Transitional woodland shrub; Coniferous forest; Natural grasslands; Mixed forest; Annual crops associated with permanent crops; Airports; Pastures.

#### Estimated ecological suitability, equines and avian biodiversity

We also found a positive relationship between equine density and estimated WNV ecological suitability (**Figure ST5**). This observation is not surprising considering that WNV historical evidence predominantly occurred in equines due to biased passive surveillance. However, as discussed in the main text, it is worth noting that as a predictor variable, the distribution of equines ranked 14 in the RF model, and 10 in the BRT model (main text **Figure 2**). That is, while a positive association between equines and suitability was recovered by the machine learning approaches, 13 and 9 other variables were deemed as more informative by the RF and BRT models, respectively. Critically, these other more relevant variables included climate, avian biodiversity and land type factors, which in contrast to equines are known, fundamental components of WNV’s zoonotic transmission cycle ^4^. As such, while we can not ignore a possible biasing of our ecological suitability towards measuring risk of spillover to equines, the predominance of other predictor variables is reassuring that a broader more informative signal is captured by the estimated ecological suitability across the country.

**Figure ST5.**
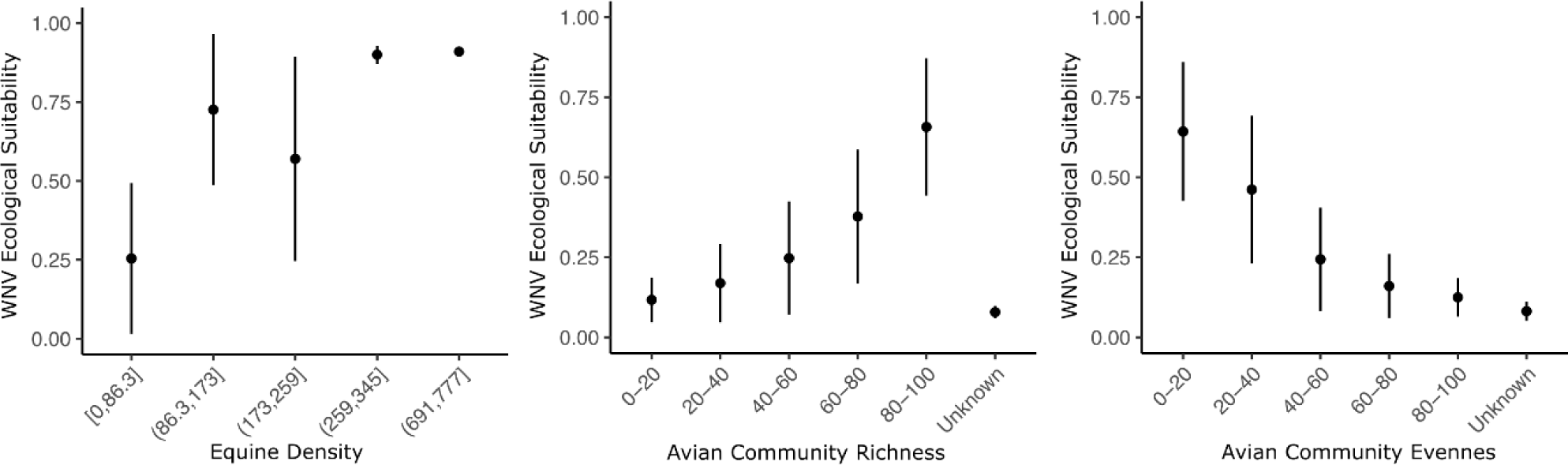
Partial plots of WNV ecological suitability in relation to equine density and avian community biodiversity metrics (richness and evenness) ranges. From left to right are equine density, avian species richness and avian species evenness. Equine density and avian species richness have a positive relation with WNV ecological suitability as opposed to avian species evenness.

Additionally, as detailed in the main text, a positive correlation was observed between suitability and avian species richness (**Figure ST5**). Species evenness is a measure that quantifies how well-balanced the frequencies of all present species are, typically presenting a negative relationship with richness (as the more species are present, the less likely it is that their frequencies are purely balanced).

#### Estimated ecological suitability and specific avian groups, families and species

We also explored the partial plots related to avian species selected as important for model prediction (by the recursive feature elimination process detailed in the **main text**). A higher relative frequency of any species implies populational dominance and consequently a reduced number of avian species (less richness) present in a specific area. We thus found that generally, an increase in the relative frequency of some specific species was associated with a decrease in WNV ecological suitability (**Figure ST6**). This was recovered e.g. for the species *Columba palumbus* (Common Wood Pigeon), *Cuculus canorus* (Common Cuckoo), *Erithacus rubecula* (European Robin), *Pica pica* (Eurasian Magpie), *Turdus merula* (Common Blackbird).

For some species, however, this logical negative relationship with the species’ relative prevalence was different. We found that while the absence of some species was associated with the lowest suitability possible, any level of relative prevalence beyond absence of those species was sufficient to guarantee a particular, non-low level of suitability (**Figure ST6**). This was recovered e.g. for the species *Anas platyrhynchos* (Mallard), *Ciconia ciconia* (White Stork), and *Hirundo rustica* (Barn Swallow). In effect, these results suggested that the presence-absence, rather than the relative prevalence of these species, was informative towards explaining the historical evidence of WNV circulation in Portugal.

**Figure ST6.**
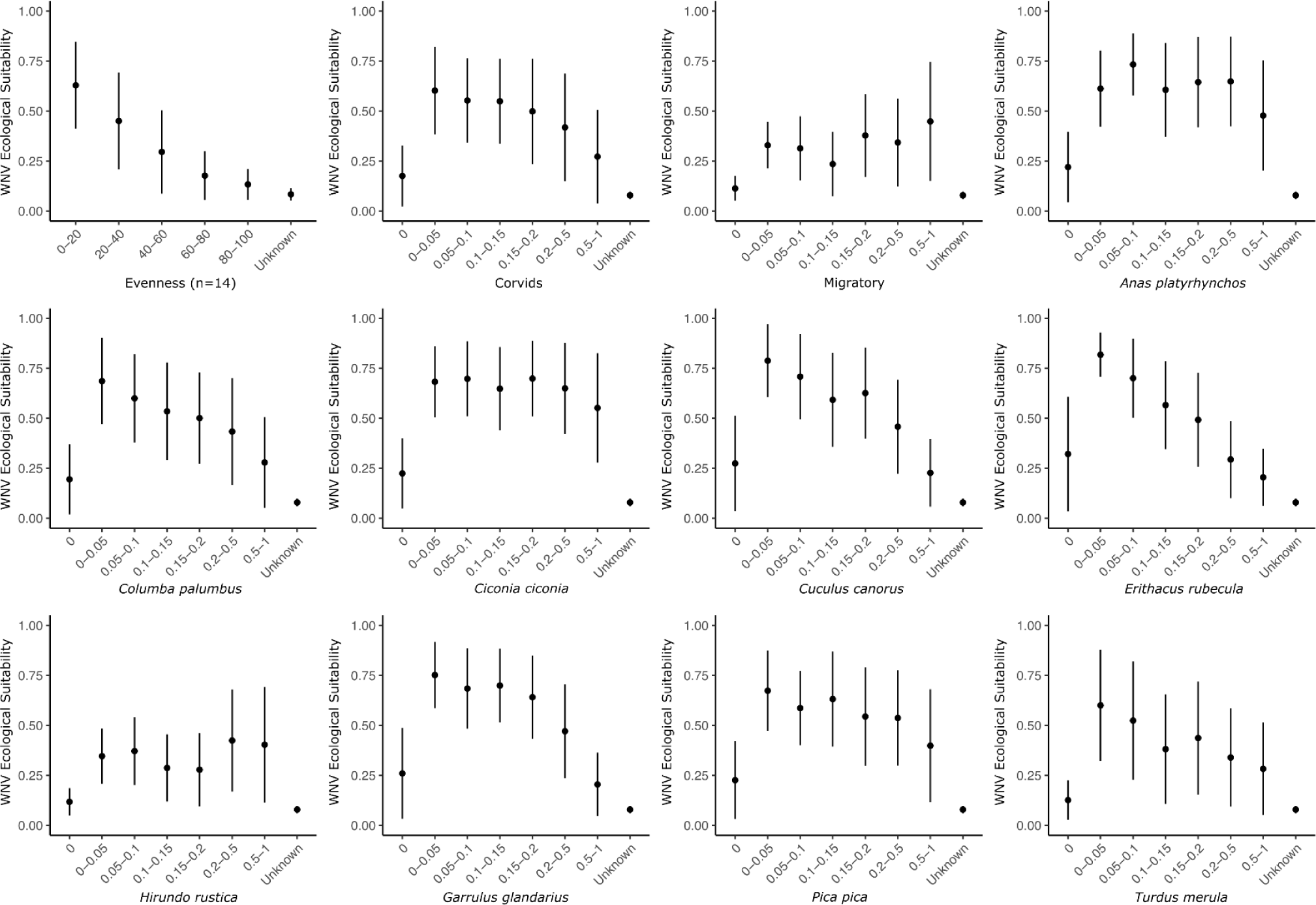
Partial plots of WNV ecological suitability in relation to avian hosts. Partial plots of evenness when considering just the 14 species selected due to their importance in WNV transmission, corvids, migratory birds and several individual species.

## Supplementary Figures

**Figure S1.**
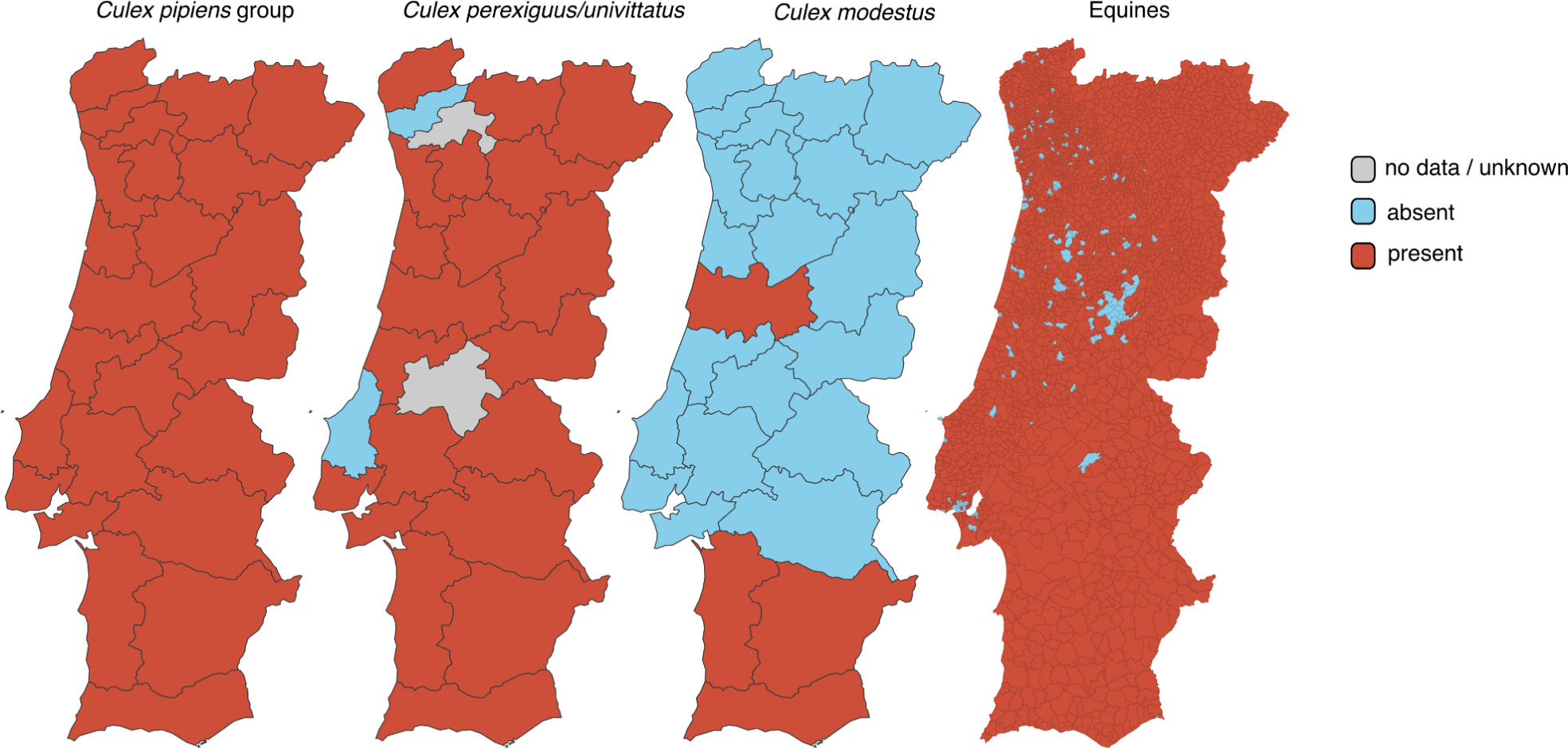
Spatial distribution of Culex spp and equines across Portugal. (3 left panels) In order, *Culex pipiens group, Culex perexiguus/univittatus, Culex modestys* and **(right)** equines. Maps related to mosquitoes use NUTS III spatial boundaries and have been replicated from the European Centre for Disease Prevention and Control (ECDC) website, last updated on October 2023 *. * https://www.ecdc.europa.eu/en/disease-vectors/surveillance-and-disease-data/mosquito-maps.

**Figure S2.**
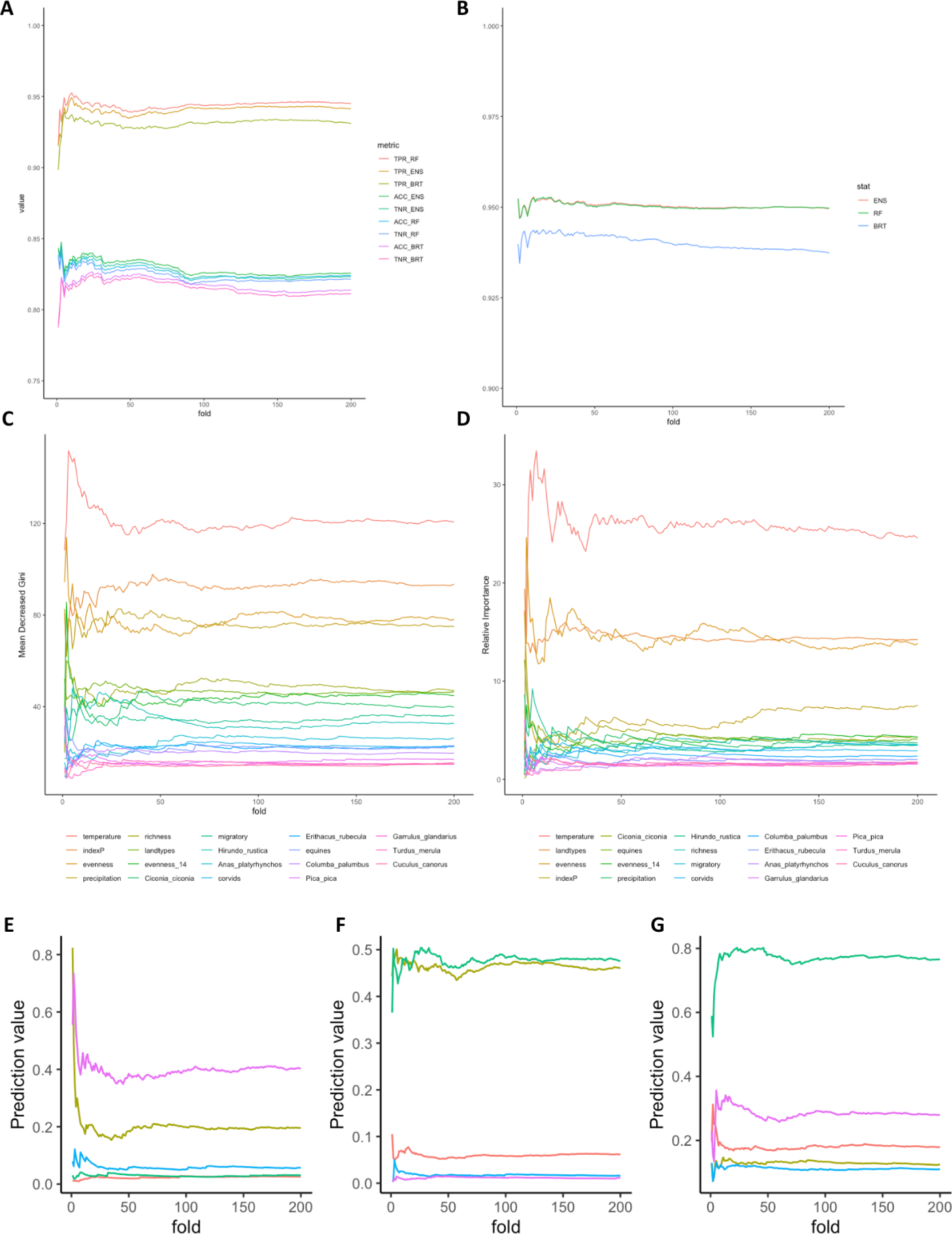
Performance metrics of all models using 0 to 200 m-folds. **(A)** Quality metrics: TPR (true positive rate), TNR (true negative rate) and accuracy for RF (random forest), BRT (boosted-regression trees) and ensemble (EM) models. **(B)** ROC for the RF and BRT models. Predictors importance estimated by **(C)** Mean Decreased Gini for RF and by **(D)** Relative Importance for BRT. **(E-G)** Prediction response stability using 0 to 200 m-folds for **(E)** BRT, **(F)** RF and **(G)** EM models (sensitivity outputs for 5 random freguesias, each with a different color).

**Figure S3.**
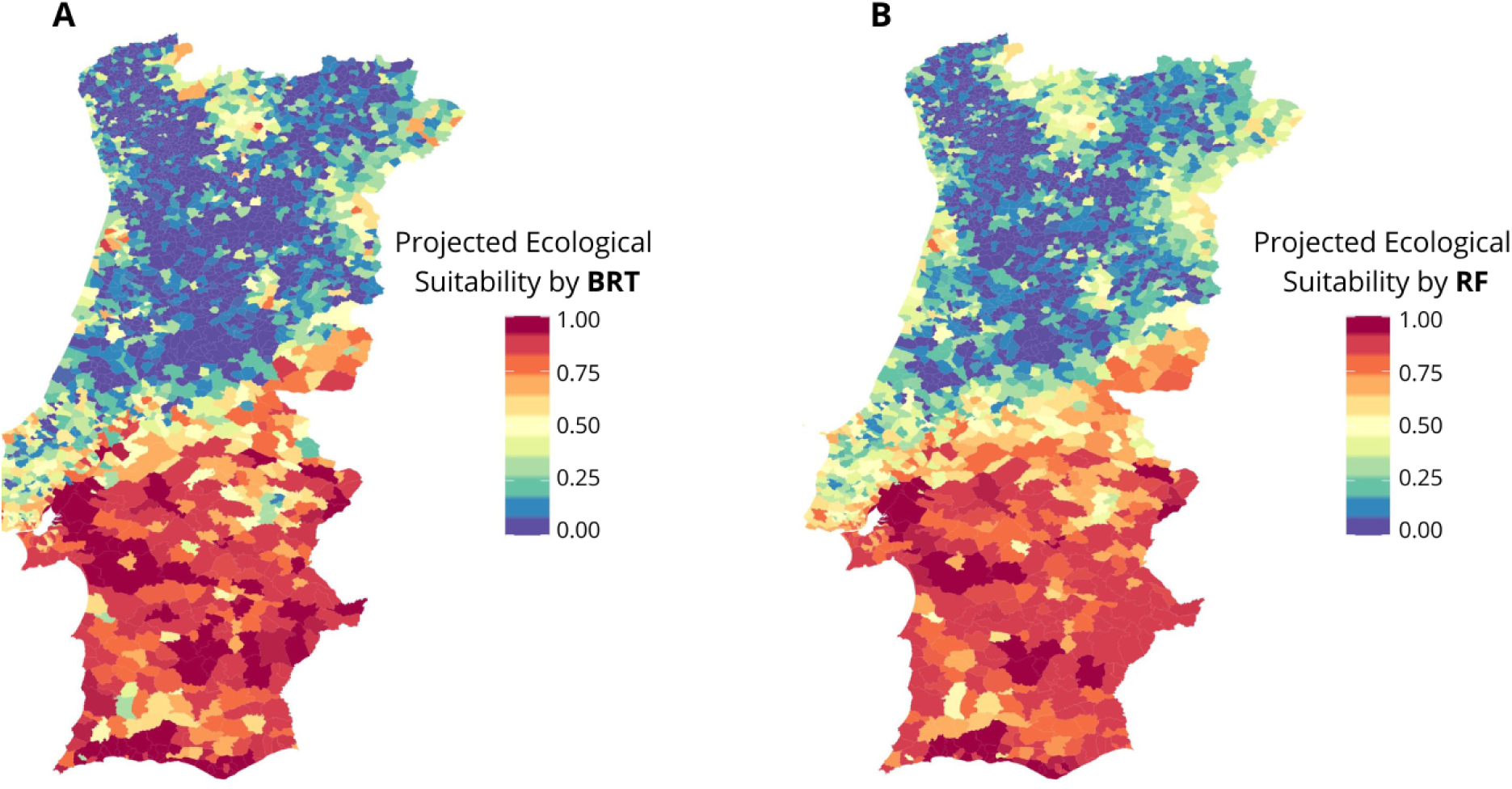
Estimated WNV ecological suitability per freguesia. **(A)** Suitability projected by the BRT (boosted-regression trees) and **(B)** RF (random forest) models, independently, per freguesia as in **Figure 3** of the main text (that presents the results from the ensemble model only).

**Figure S4.**
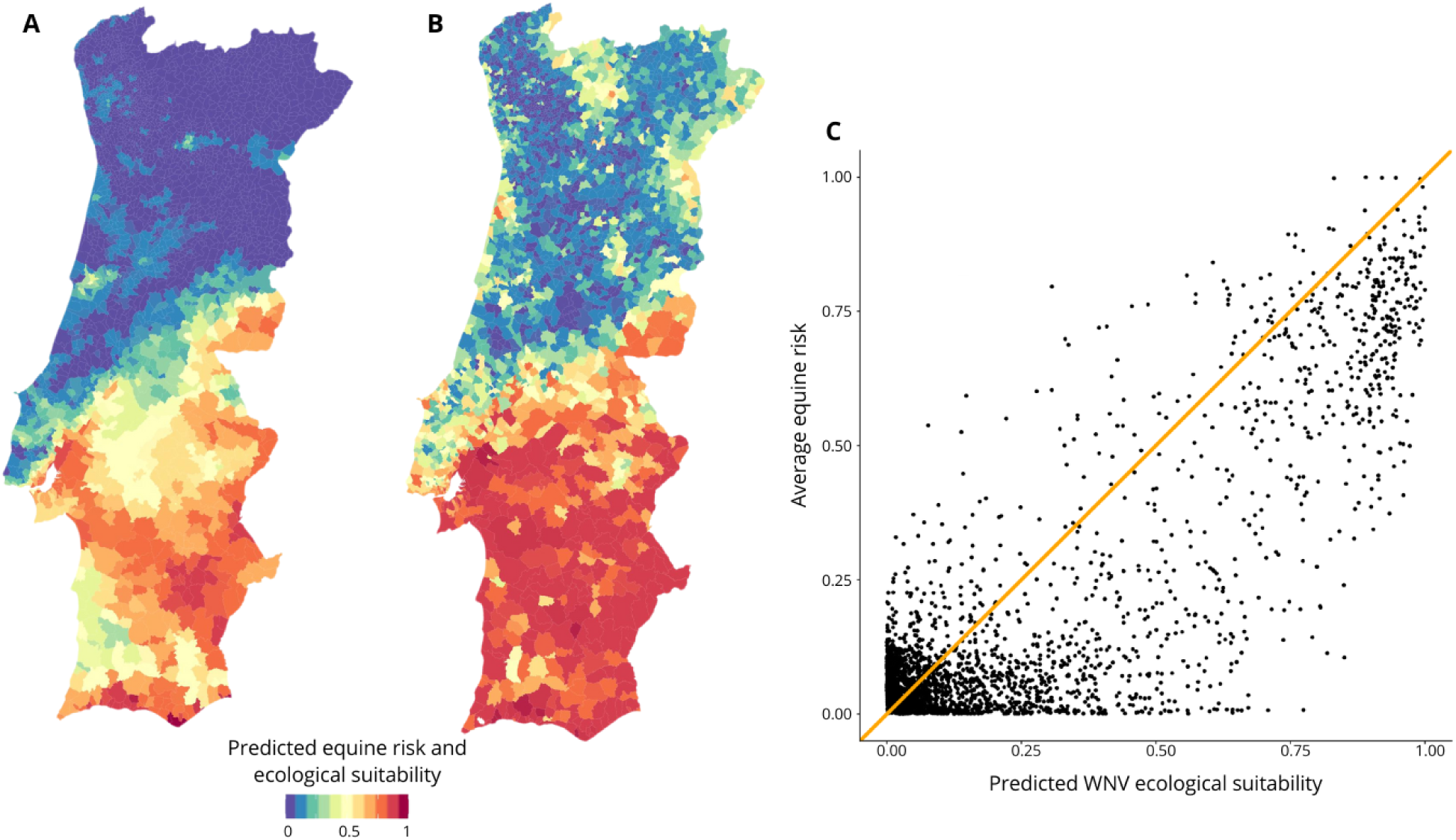
Comparison of modelling outputs from this study with modelling outputs from a recent study at the Iberian peninsula level. **(A)** Estimated WNV equine risk map for Portugal was extracted from published and shared results of the study by García-Carrasco and colleagues ^1^. **(B)** Estimated WNV risk map for Portugal from this study (Figure 3, main text). **(C)** Comparison of suitability per freguesia between this study and García-Carrasco, J.-M. et al. West Nile virus in the Iberian Peninsula: using equine cases to identify high-risk areas for humans. Eurosurveillance 28, 2200844 (2023).

**Figure S5.**
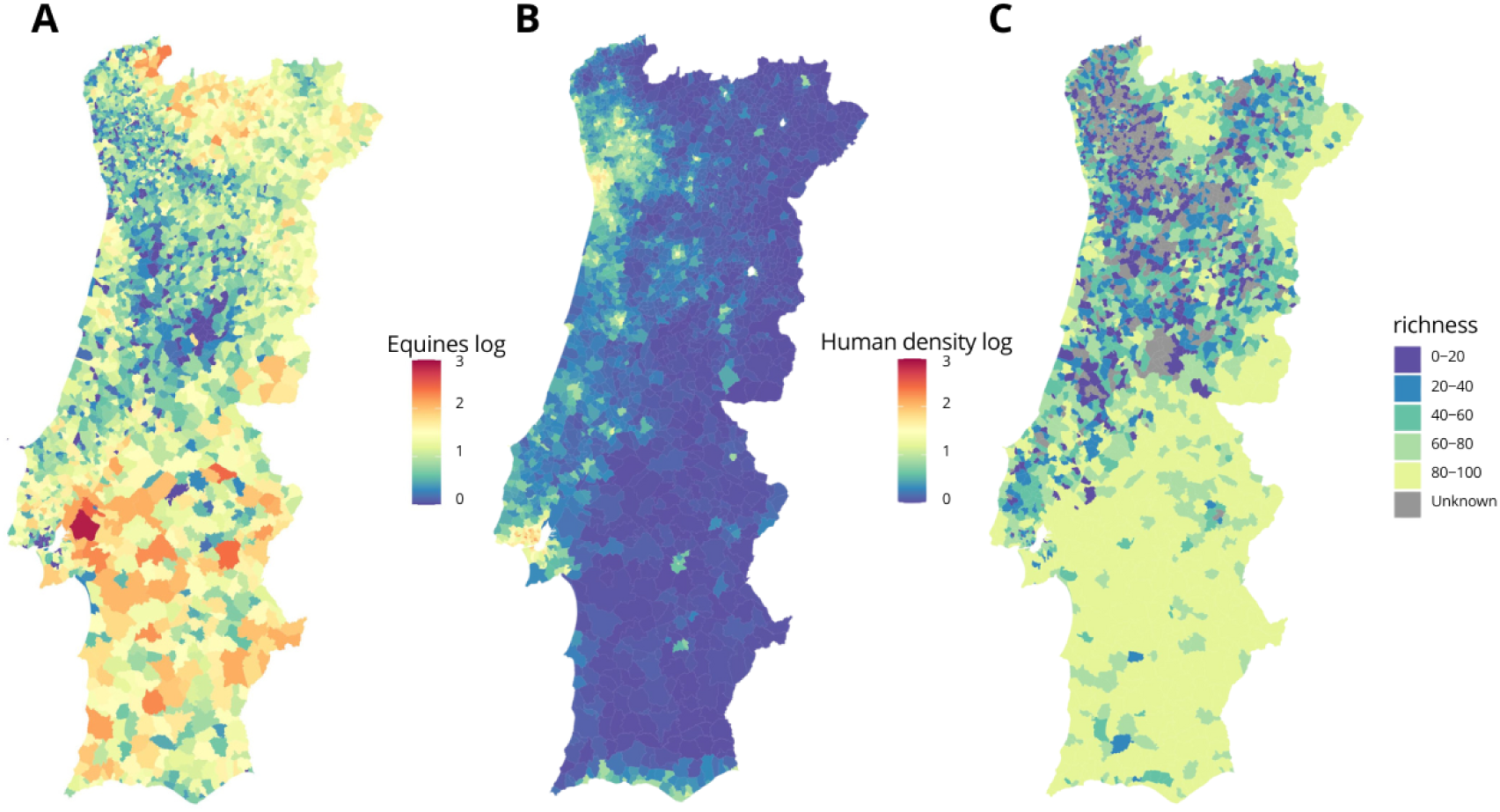
Spatial distributions of 3-host axes. **(A)** Equines (log of equine number). **(B)** Human density (log of density). **(C)** Avian biodiversity (species richness), where grey marks freguesias with missing data.

**Figure S6.**
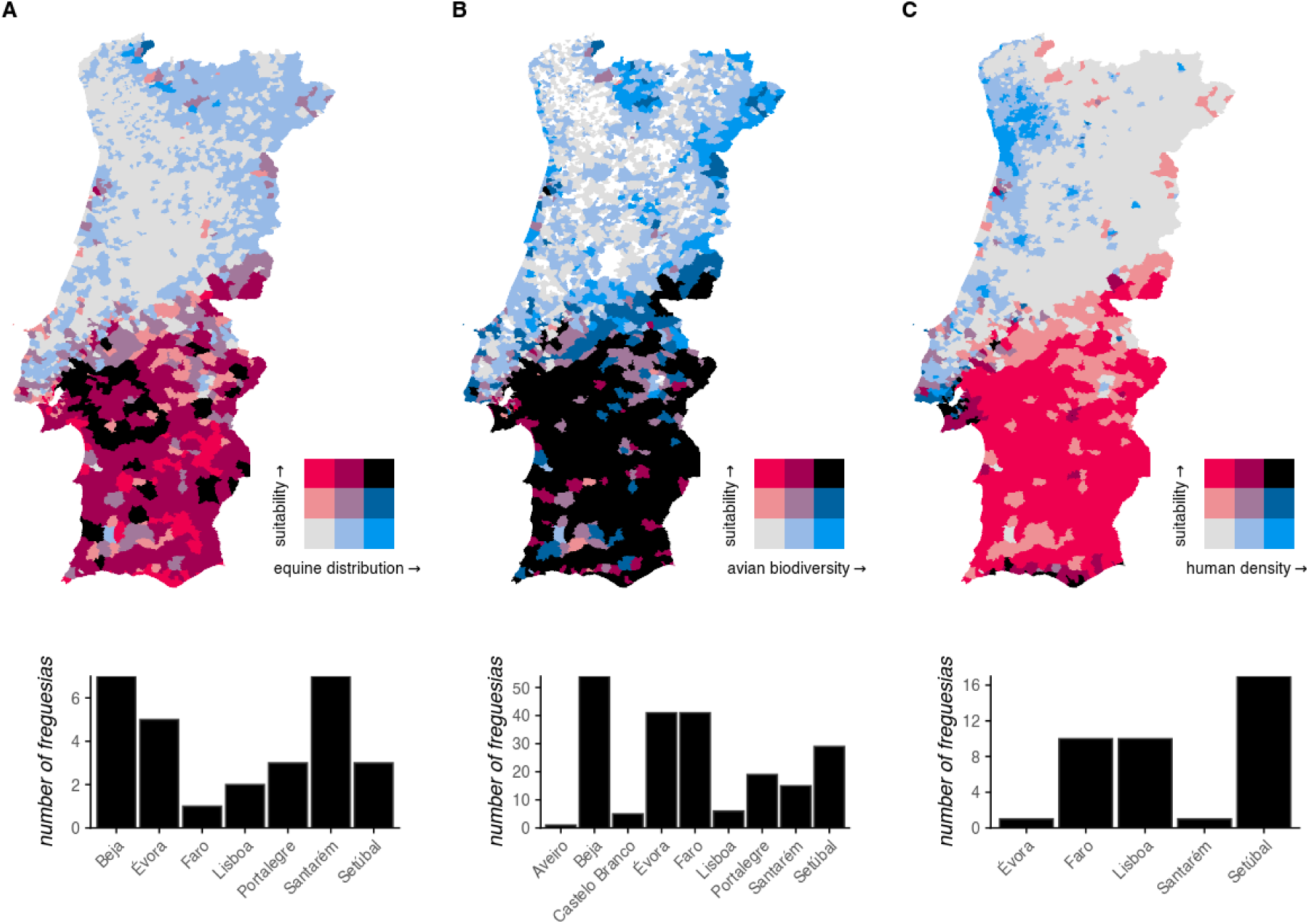
Bivariate distributions of suitability versus 3-host axes. (left) Suitability vs equines (absolute numbers). (**center**) Suitability vs avian biodiversity (richness), where white freguesias mark missing data. (**right**) Suitability versus human density. Three classes per variable were considered in order to obtain a 3×3 bivariate color scale: suitability as in the categories A-C from main text; equine distribution 0-10, 10-100, +100; human density 0-1, 1-5, +5. **Top panels** are the bivariate distributions. **Bottom panels** are total freguesias per district (x-axis) that feature combinations of high suitability with each of host axes (black color on the maps)

**Figure S7.**
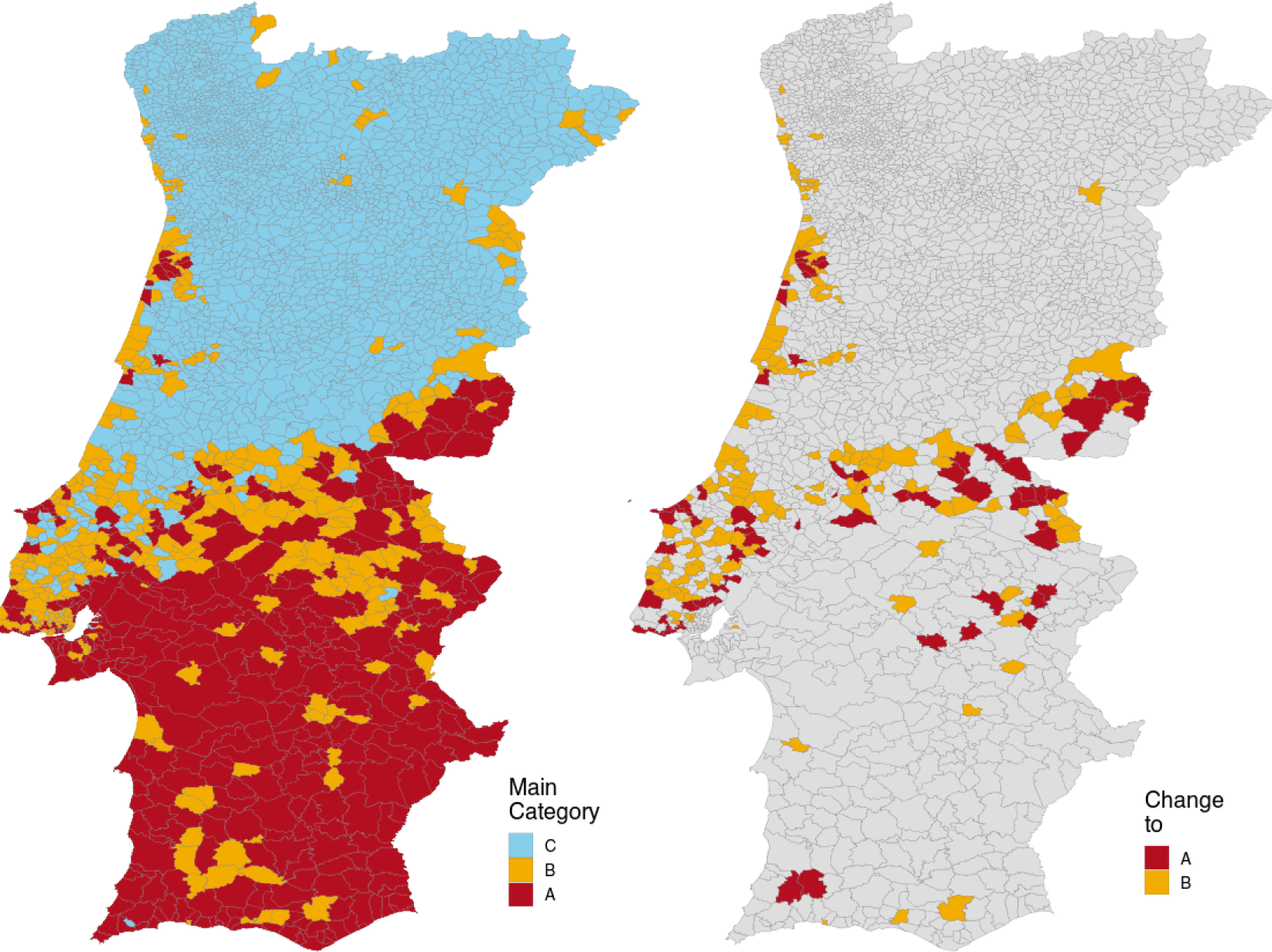
Main categories per freguesias resulting from projected effects of climate change. (left) Map showing the distribution of the 3 main categories in the year 2050 (to be compared with main **Figure 4** for 2022). **(right)** Highlighted are freguesias that change from categories C and B to A, or from C to B, when assessing their current category in 2022 (Figure 4 main text) and their projected category in 2050 (left panel of this figure). Those not changing category are presented in grey.

